# Tissue homeostasis and adaptation to immune challenge resolved by fibroblast network mechanics

**DOI:** 10.1101/2021.05.27.446027

**Authors:** Harry L. Horsnell, Robert J. Tetley, Henry De Belly, Spyridon Makris, Lindsey J. Millward, Agnesska C. Benjamin, Charlotte M de Winde, Ewa K. Paluch, Yanlan Mao, Sophie E. Acton

## Abstract

Emergent physical properties of tissues are not readily understood by reductionist studies of their constituent cells. Here, we show molecular signals controlling cellular physical properties, collectively determining tissue mechanics of lymph nodes, an immunologically-relevant, adult mammalian tissue. Lymph nodes paradoxically maintain robust tissue architecture in homeostasis yet are continually poised for extensive tissue expansion upon immune challenge. We find that following immune challenge, cytoskeletal mechanics of a cellular meshwork of fibroblasts determine tissue tension independently of extracellular matrix scaffolds. We determine that CLEC-2/podoplanin signalling regulates the cell surface mechanics of fibroblasts, permitting cell elongation and interdigitation through expedited access to plasma membrane reservoirs. Increased tissue tension through the stromal meshwork gates the initiation of fibroblast proliferation, restoring homeostatic cellular ratios and tissue structure through expansion.

---

Unlike developmental systems that display progressive tissue growth and maturation^1^, the homeostatic state of adult tissues is robust, maintaining form and function^2^. Secondary lymphoid organs, are uniquely able to dramatically change size in response to immune challenge, adapting to the increased space requirements of millions of infiltrating and proliferating lymphocytes, while remaining structurally and functionally intact throughout^3–5^. As a mechanical system, lymph nodes continually resist and buffer the forces exerted by trafficking of lymphocytes entering and leaving the tissue in steady state^6^, and managing diurnal fluctuations in cell trafficking^7^. Tissue size is determined by lymphocyte numbers, highlighted by the small organ size in genetic models blocking lymphocyte development (Rag1 KO)^8^, and the ability of the tissue to expand 2-10 fold in size to accommodate lymphocyte proliferation through adaptive immune responses^3–5,9^.

Lymph nodes function at the interface of immunity and fluid homeostasis, constructing a physical 3-dimensional cellular meshwork linking fluid flow, immune surveillance and adaptive immunity^10^. The most populous stromal cell component are fibroblastic reticular cells (FRCs) which span the whole tissue generating a interconnected cellular network with small world properties^11^, forming robust clustered nodes with short path lengths, and surrounding bundles of extracellular matrix fibres^12^. It is widely assumed that extracellular matrix scaffolds are the predominant force-bearing structures determining tissue mechanics^1,2^. However the relative contributions of the cellular structures versus the underlying extracellular network to tissue mechanics have not been addressed in a highly cellularised system undergoing such extensive expansion^2^. As tissue size and cellularity increase in response to immunogenic challenge, it is known that lymph nodes become more deformable when challenged with compressive force^3^. During the remodelling process the FRC network maintains connectivity through the elongation and increased spacing between FRCs, increasing mesh size of the network^3^. It is also known that CLEC-2^+^ dendritic cells are required to prime the stromal architecture for tissue expansion^3,4^ but the downstream impacts on the mechanical properties of the cellular network and extracellular matrix scaffolds driving the adaptation in tissue mechanics are unknown. During tissue expansion, FRCs reduce their adhesion to the underlying extracellular matrix bundles and these matrix scaffolds become fragmented^9^. This makes lymph nodes an ideal model system to address the relative mechanical contributions of cellular and material structures to emergent tissue mechanics.

As fibroblasts are contractile, force-generating cells^13^, we hypothesise that the interconnected FRC network determines tissue mechanics. The FRC network, identified by podoplanin expression, spans the whole lymph node tissue but specifically supports CD3^+^ T cell function in the paracortex, providing trafficking routes from high endothelial venules to B cell follicles^14,15,16,17^ (Fig. 1A-B). We asked how the FRC network and associated extracellular matrix scaffolds participate in tissue mechanics and architecture in the immunological steady state. Following laser ablation (Fig. 1C-D, and supplementary Movie. S1A), we measured a mean recoil of 0.42*μ*m/s^-1^ in the fibroblastic reticular network of the paracortex, using a fibroblast-specific membrane-eGFP mouse model (PDGFR*α*-mGFP-CreERT2 (Fig. 1C, Fig. S1, Fig. S2A), formally demonstrating that the reticular network is under mechanical tension in the tissue (Fig. 1E).

**Figure 1 –.**
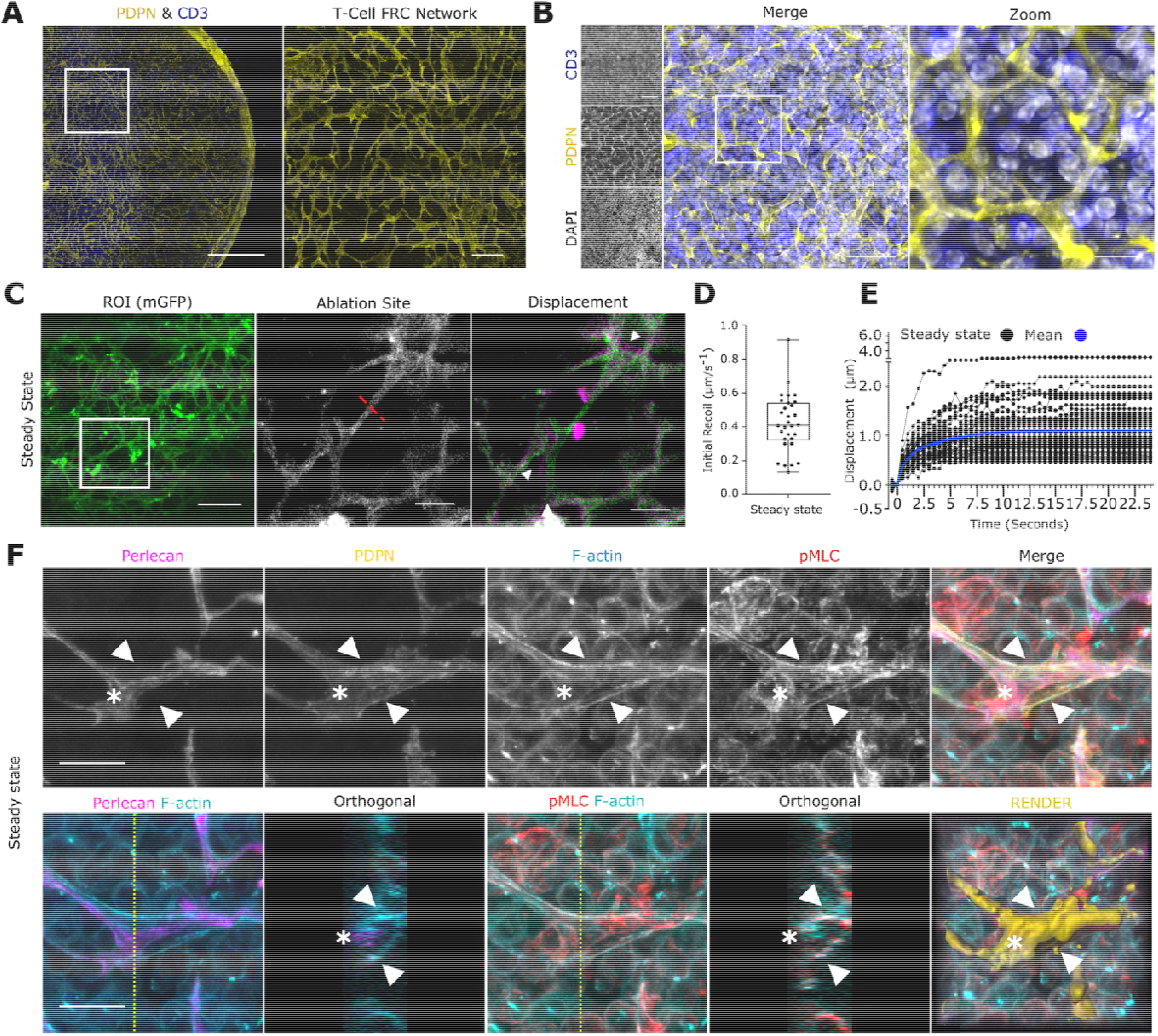
Lymph node tissue tension measured through the FRC network. **A)** Lymph node tile scan (left), paracortex (right) maximum Z-projection. PDPN (FRCs, yellow), CD3 (T-cells, blue). Scale bar 500 m (left), 25 m (right). **B)** FRC network structure and T-cell compaction vibratome slices. PDPN (FRCs), CD3 (T-cells), DAPI (Nuclei). Scale bar (left), (right). **C)** PDGFR -mGFP (FRCs) ablation ROI (white box) and cut site (red dotted line). Scale bar. Recoil displacement (white arrowheads) with pre (green) and post cut (magenta) overlay. Scale bar **D)** Initial recoil velocity (m/s^-1^). Box plot indicates median, interquartile range and min/max. **E)** Individual recoil curves (black) compared to the mean (blue). N=5 LNs. F) Paracortical steady state FRCs. Perlecan (matrix), PDPN (FRC), phalloidin (F-actin) and pMLC. Asterisk and arrowheads indicate F-actin cables. Orthogonal views (yellow dotted line, YZ axis, 10*μ*m depth). Scale bar 10*μ*m.

To investigate how cellular-scale components of the stromal architecture contribute to tissue mechanics, we examined the cytoskeletal and extracellular matrix structures of the reticular network. In steady state, FRCs tightly adhere to and enwrap collagen bundles forming a continuous meshwork^18^. Therefore, recoil following laser ablation is a measure of the combined mechanical properties of both the cell and the extracellular matrix structures in steady state (Fig. 1C-F). Maximum z-projections and orthogonal views of FRCs show F-actin cables aligned proximal to the collagen bundles and spanning beneath the T-cell-facing FRC plasma membrane (Fig. 1F, and supplementary Movie. S1B). These F-actin structures co-localised with phosphorylated myosin regulatory light chain (pMLC2)^19^ (Fig. 1F), indicating that FRCs are contractile and resisting strain in steady state^20^. Since resting lymph node size is constant, forces are balanced in the steady state tissue.

We next asked how the reticular network reacts to the forces exerted by increasing numbers of lymphocytes following immune challenge. It has been observed that the network of bundled matrix fibres becomes fragmented^9^, while the cellular component of the meshwork remains intact and connected^3,4,11^. Therefore, we next tested if the intact cellular network could compensate for the loss of extracellular matrix integrity to balance tissue tension. In response to immunisation with incomplete Freund’s adjuvant and ovalbumin (IFA/OVA), the tissue expands 2-3 fold over the first 5 days (Fig. 2A). Surprisingly, at day 3 post-immunisation, initial recoil velocity was decreased by 29% to 0.30μm/s^-1^ (Fig. 2B-D, and Supplementary Movie. S2A) despite a 1.5-fold increase in tissue mass (Fig 2A). In contrast, 2 days later (at day 5 post-immunisation), mean initial recoil velocity was 60% higher than in the steady state at 0.71μm/s^-1^ (Fig. 2B-D, and supplementary Movie. 2B). Since the extracellular matrix scaffold is fragmented at this phase of tissue expansion^9^, we hypothesised that the actin cytoskeleton must be the major contributor to the increased tissue tension. Indeed, recoil measurements following laser ablation of tissues pre-treated with ROCK inhibitor (Y27632) to inhibit cytoskeletal contractility^19^, demonstrated that tissue tension was reduced to basal levels at all time points tested (Fig. 2C, Fig. S2B-F). These results indicate that tissue tension is determined by differential resistive actomyosin forces (Fig.2C).

**Figure 2 –.**
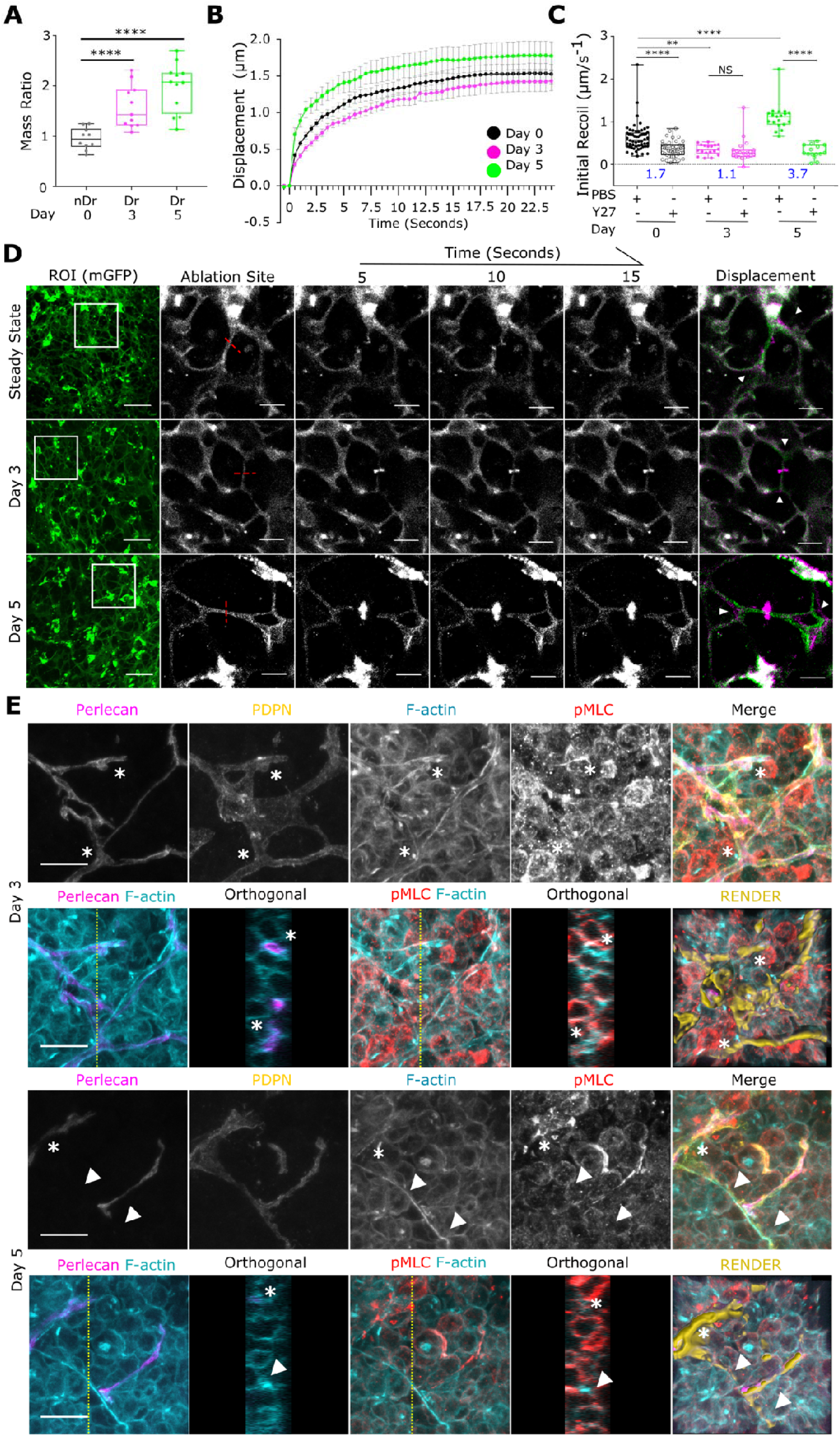
Actomyosin contractility sets FRC network tension in response to tissue expansion. **A)** Lymph node Mass post IFA/OVA immunisation. N>5 **B)** Recoil curves of network displacement (*μ*m) (mean ±SEM). N>5 **C)** Initial recoil velocity (*μ*m/s^-1^) post IFA/OVA immunisation, +/- ROCK inhibition (Y27632). Resistive myosin forces (blue text = ratio between PBS/Y27). Box plot indicates median, interquartile range and min/max. Two-way ANOVA with Tukeys multiple comparisons, **p<0.01, ****p<0.0001. N>5. **D)** PDGFR*α*-mGFP (FRCs) ablation ROI (white box) and cut site (red dotted line). Scale bar 50**μ*m*. Recoil displacement (white arrowheads) with pre (green) and post cut (magenta) overlay. Scale bar 10**μ*m*. **E)** Paracortical FRCs post IFA/OVA immunisation. Perlecan (matrix), PDPN (FRC), phalloidin (F-actin) and pMLC. Asterisk and arrowheads indicate F-actin cables. Orthogonal views (yellow dotted line, YZ axis, 10*μ*m depth). Scale bar 10*μ*m.

5 days post immunisation, the whole tissue is more deformable^3^, yet tension measured through the fibroblastic reticular network increases (Fig. 2B-C). Since elasticity of gels is known to scale as the inverse of mesh size^21^, the combination of the fragmentation of the matrix and increasing mesh size of the FRC network may explain increased tissue deformability through acute expansion. To investigate how the FRC network responds mechanically to the increasing tissue size (Fig. 2A) we compared the cytoskeletal and matrix structures of the FRCs at day 3 (lower tension) versus day 5 (higher tension). In reactive lymph nodes, 3 days after IFA/OVA immunisation, F-actin structures were located primarily proximal to the basement membrane (asterisk), but T-cell facing F-actin cables were absent (Fig. 2E, and Supplementary Movie S2C). However, 5 days post immunisation, prominent contractile F-actin cables structures span the length of FRC cell bodies even in the absence of underlying matrix bundles (arrowhead) (Fig. 2E, and supplementary Movie S2D). These data show that tissue tension varies in response to immunogenic challenge and that increased tissue tension occurs independently of extracellular matrix integrity. Instead, mechanical forces are generated by increased packing of lymphocytes in the FRC meshwork and resisted by actomyosin through the FRC network (Fig. 2).

Since the FRC network is not resisting increased packing of lymphocytes at day 3 (Fig. 2C), regulation of actomyosin contractility alone cannot explain how the FRC network remains connected as the tissue mass increases (Fig. 2A). We next sought to understand the mechanisms controlling the cell surface mechanics of FRCs. To increase mesh size without increasing FRC number^3,4^ FRCs must elongate or make protrusions, adapting their cell morphology to maintain network integrity^3,11^. Effective membrane tension (hereafter membrane tension), is determined by the in-plane tension of the lipid bilayer and the strength of membrane-to-cortex attachments^22,23^ and can regulate cell spreading, changes in morphology and cell fate^22–24^. Cell contact between antigen-presenting dendritic cells and FRCs through CLEC-2/podoplanin binding, peaking at day 3, is known to regulate lymph node deformability and tissue expansion^3,4^. Therefore, we asked whether the CLEC2/podoplanin interaction regulates FRC cell surface mechanics. We used optical tweezers to measure membrane tension of FRCs, (Fig. 3A, Fig. S3G, and supplementary Movie. S3A) and find that specific engagement of CLEC-2 to podoplanin reduces membrane tension (Fig. 3A-B) and downregulates phosphorylation of ezrin, radixin and moesin family proteins (pERM) (Fig. S3F), which tether the actin cytoskeleton to the plasma membrane. Signalling specificity was confirmed by exogenous expression of a PDPN mutant that cannot bind CLEC-2 (T34A) and a mutant which cannot signal through the cytoplasmic tail (S167A-S171A) (Fig. S3A-D). Both showed no change in membrane tension in the presence of CLEC-2 (Fig. S3A-D). As podoplanin interacts with CD44^25^, also known to bind ezrin^26^, we tested the relatively contributions of both CD44 and podoplanin to membrane tension in unstimulated FRCs and find that podoplanin is the key driver in steady state (Fig. 3C, Fig. S3E)

**Figure 3 –.**
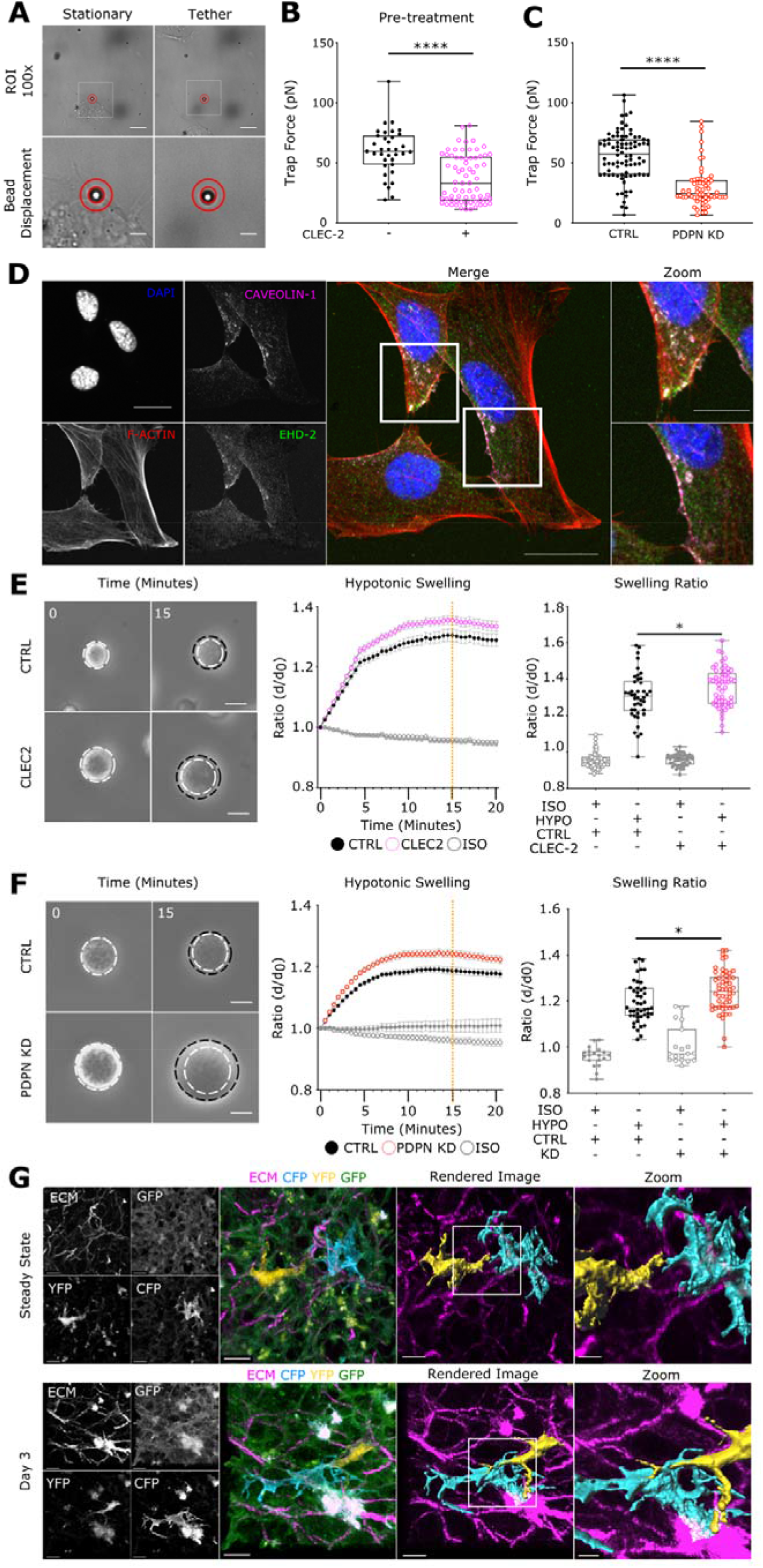
CLEC-2/PDPN control intrinsic FRC mechanics for morphological adaption. **A)** Optical tweezers generate membrane tethers of FRCs, scale bar 10*μ*m. Stationery (bottom left) and tether (bottom right) bead displacement (red circles). Scale bar 2*μ*m. **B)** Trap force (pN/*μ*m) of FRCs pre-treated with CLEC-2. Box plot indicates median, interquartile range and min/max. Mann-Whitney test, p<0.001. Points represent individual cells. N>35. **C)** Trap force (pN/*μ*m) of PDPN shRNA KD FRCs Trap force measurement of FRCs. Box plot indicates median, interquartile range and min/max. Mann-Whitney test, ****p<0.001. Points represent individual cells. N>64. **D)** Caveolae structures in FRCs, scale bar 20*μ*m. DAPI (Blue), F-actin (Red), Caveolin-1 (magenta) and EHD2 (Green). Zooms (white box), scale bar 10*μ*m. **E)** Swelling of PDPN CTRL FRCs +/- pre-treatment CLEC2. Initial size (white circle) post swelling (black circle). Scale bar 25*μ*m. Change in diameter ratio (middle, mean ±SEM). Diameter ratio of control and CLEC2 treatment at 15 minutes post swelling (orange dotted line, (right)). Box plot indicates median, interquartile range and min/max. One-way ANOVA with Tukeys multiple comparisons, p<0.01. N>42. **F)** Swelling of PDPN shRNA KD FRCs. Initial size (white circle) post swelling (black circle). Scale bar 25*μ*m. Change in diameter ratio (middle, mean ±SEM). Diameter ratio of control and CLEC2 treatment at 15 minutes post swelling (orange dotted line, (right)). Box plot indicates median, interquartile range and min/max. One-way ANOVA with Tukeys multiple comparisons, *p<0.01. N>19. **G)** Individually labelled FRCs *in vivo*, scale bar 10*μ*m. Zoomed region (white box), scale bar 5*μ*m.

In the tissue, FRCs must each elongate and form protrusions to interdigitate with their network neighbours as they spread and expand the stromal cell network. These cell shape changes can be achieved by increasing the ratio of plasma membrane surface area to cell volume through exocytotic pathways or by unfolding membrane reservoirs^27^. FRCs contain active EHD2^+^caveolae structures^28^ contributing to tension-sensitive plasma membrane reservoirs (Fig. 3D). We tested, using osmotic shock (Fig. S4A-B) whether regulation of membrane tension through the CLEC-2/podoplanin signalling axis impacted how rapidly FRC could utilise existing membrane reservoirs to respond to the external forces (Fig 3E-F). We found that CLEC-2 engagement to podoplanin (Fig. 3E, Fig. S4C, and supplementary Movie. S4-B) or knockdown of podoplanin (PDPN KD) (Fig. 3F, and supplementary Movie. S4C) permitted more rapid cell expansion in hypotonic conditions but did not alter total membrane availability (Fig. S4D, and supplementary Movie. S4D). *In vivo*, we observed individual, differentially labelled, neighbouring FRCs increasing cell-cell contact 3 days post immunisation (Fig. 3G), suggesting that additional plasma membrane is used for morphological adaptations and incorporated into cell extensions. We propose that changing cell surface mechanics of FRCs in combination with reduced actomyosin resistance (Fig. 2C) permits lymph node expansion whilst maintaining FRC network integrity.

However, at day 5, even larger number of lymphocytes puts the network under increased tension which is resisted by actomyosin contractility (Fig. 2C). We asked what the functional consequence of changing tissue tension had on tissue remodelling in response to immune challenge. Existing studies have found that expansion of FRC populations lag behind lymphocytes^3–5^, but it is not known how FRC proliferation is triggered or spatially regulated within the tissue. 5 days post IFA/OVA immunisation the number of proliferating FRCs (Ki67^+^) doubles compared to steady state (Fig. 4A-B), which correlates with increased tissue tension (Fig. 2B-D). Ki67^+^ proliferative FRCs were observed throughout the tissue, and we observed no specific proliferative niche surrounding blood vessels or beneath the bounding capsule (Fig. 4A), suggesting that the cue for entry into the cell cycle is not spatially restricted or limited to a subpopulation of FRCs. We hypothesised that increased mechanical tension may gate FRC entry into cell division. To test this, we blocked the increase in tissue tension unilaterally *in vivo* through pharmacological inhibition of ROCK^19^ (Fig. 4C-D) and found that proliferation of stromal cells, specifically the FRC population, was significantly attenuated 5 days post immunisation, while lymphocyte proliferation and the increases in tissue mass and cellularity were unaffected (Fig. 4E-I, Fig. S5A-D). This leads us to conclude that the stromal architecture is reactive to the physical space requirements of the lymphocyte populations. Indeed, previous studies have identified a remarkably robust ratio between fibroblastic stroma and T cells, maintained as the tissue expands^29^. A mechanical cue for stromal cell growth and proliferation would ensure that the steady state ratio of fibroblastic stroma to lymphocytes is perfectly restored, independently of the kinetics or scale of the immune reaction. Homeostatic tissue architecture is recovered as the tissue expands reinstating the supportive immune microenvironment for lymphocyte populations^30^.

**Figure 4 –.**
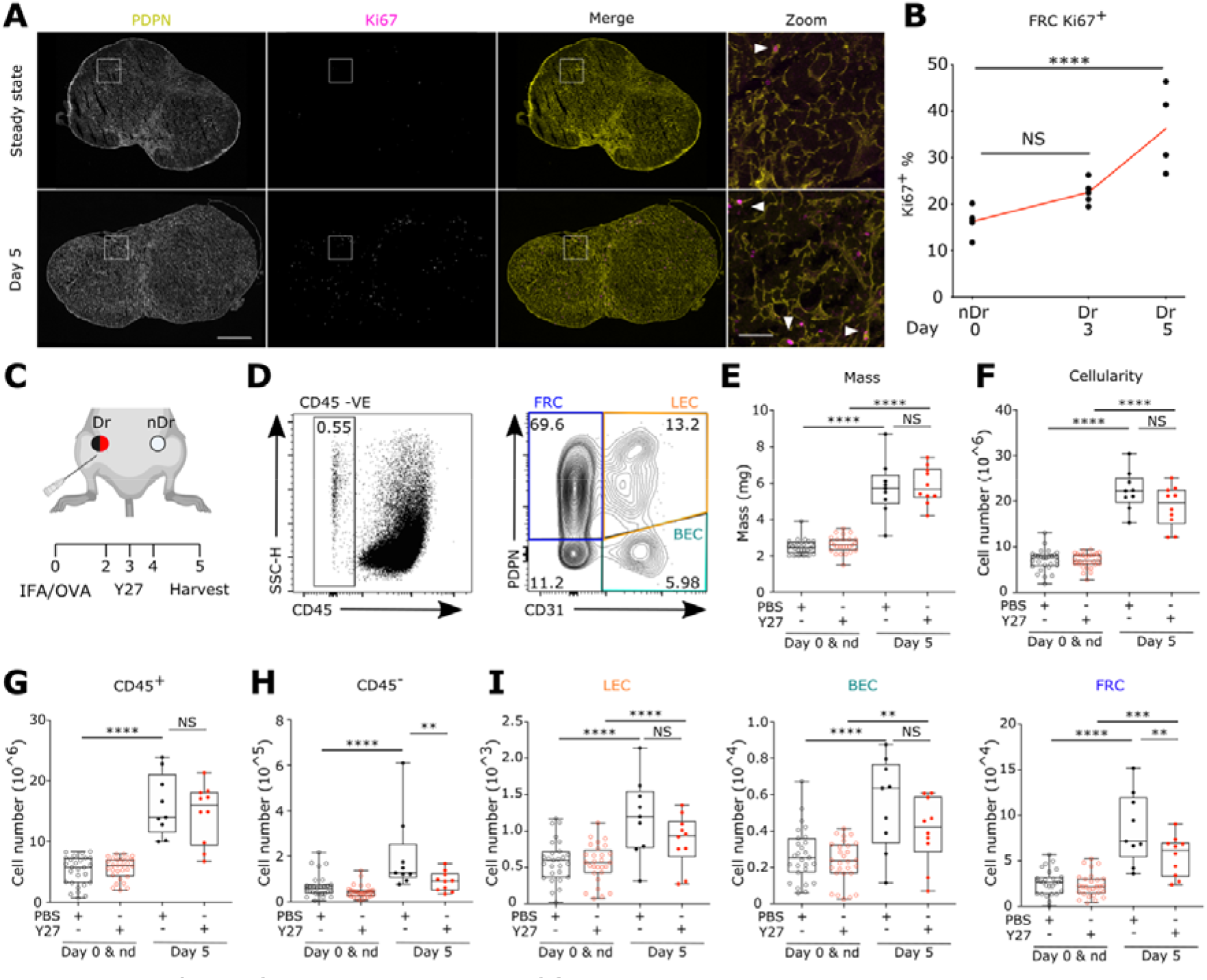
Mechanical tension gates FRC proliferation. **A)** LN tile scan of PDPN^+^ (left), Ki67^+^ (middle). Scale bar 500 m. zoom in T-cell area (white box). Scale bar 50 m. Arrowheads mark PDPN^+^ Ki67^+^ FRCs. **B)** Flow cytometric analysis of Ki67^+^ % of FRC’s post IFA/OVA immunisation. Two-way ANOVA with Tukeys multiple comparisons, ****p<0.0001. N>4. **C)** Schematic of immunisation and Y27 treatment timeline. **D)** Flow cytometric gating, representative percentages of CD45-stroma cells (left) and sub-populations (right). Fibroblastic reticular cells (FRC-PDPN^+^CD3Γ), blood endothelial cells (BEC-PDPN^-^CD31^+^), lymphatic endothelial cells (LEC-PDPN^+^CD31^+^). **E)** LN mass. Box plot indicates median, interquartile range and min/max. Two-way ANOVA with Tukeys multiple comparisons, ****p<0.0001, ***p<0.001. N>9. **F-G)** Total number of cells post IFA/OVA immunisation, +/- ROCK inhibition (Y27632). Immune cells (CD45^+^), stroma (CD45^-^). **I)** Total number of LECs (left), BECs (middle) and FRCs (right) post IFA/OVA immunisation, +/- ROCK inhibition (Y27632). Box plot indicates median, interquartile range and min/max. Two-way ANOVA with Tukeys multiple comparisons, ****p<0.0001, ***p<0.001, **p<0.01. N>9.

Together our data demonstrate that the fibroblastic structure of the lymph node is the active mechanical component during tissue expansion. Using the dynamic cellular network rather than the more rigid ECM to respond to changing lymphocytes numbers in the tissue provides the lymph node with an elegant mechanical system that can proportionately respond to lymphocyte requirements. We show that the lymph node becomes mechanically permissive to expansion through CLEC-2/PDPN signalling priming the cell surface mechanics of FRCs. As the kinetics of expansion increase tissue tension, FRC’s initiate cellular division to restore tissue architecture. Other studies have shown that tissue scale properties emerge from cellular scale mechanics in the transition from developing to adult tissue^31^. We now directly address this concept in an immunological relevant adult mammalian tissue during homeostasis and immune challenge. We show that the interconnected cellular network deploys molecular signals controlling cellular mechanical properties to collectively determine tissue scale mechanics of lymph nodes.

## Acknowledgements

We thank Prof E. Sahai, Prof G. Charras and Dr V.G. Martinez for critical reading of the manuscript. We also thank Prof. H. Clevers for supplying R26R-confetti mice.

## Author contributions

H.L.H and S.E.A designed the study and wrote the manuscript. H.L.H performed and analysed all experiments. A.C.B. and S.E.A. generated the PDGFR*α*-mGFP-CreERT2 mouse model. R.J.T assisted with laser ablation studies. H.d.B assisted with optical trap measurements. H.d.B contribution was carried out under the support of E.K.P at MRC-LMCB UCL, current location is the department of Biochemistry and Biophysics, UCSF, San Francisco, CA, USA. S.M assisted with flow cytometry of stromal cells. L.J.M conducted imaging experiments of FRCs *in vivo*. C.M.d.W. generated CD44 KO cell lines. Y.M and E.K.P contributed to study design and editing of the manuscript. All authors contributed to editing of the manuscript.

## Supplementary Materials

### Materials & Methods

#### Mice

Experiments were performed in accordance with national and institutional guidelines for animal care and approved by the institutional animal ethics committee review board, European research council and the UK Home office. Breeding of animal lines were maintained off site at Charles River Laboratory. Wild type C57BL/6J mice were purchased from Charles River Laboratories. Novel mouse model of PDGFR*α*-mGFP-CreERT2 generation. EGFP with a membrane tag (N-terminal 0-20 amino acids of neuromodulin GAP-43) was inserted into the PDGFR*α* gene locus in combination with a CreERT2 cassette (linked to mGFP with a P2A self cleavage peptide^1^) using Cyagen CRISPR Cas9 technology. PDGFR*α*-mGFP-CreERT2 x R26R-Confetti^2^ mice were also generated. Females and males aged between 8-12 weeks were used for experiments, unless stated otherwise.

#### Immunisation Model

Mice were immunised via subcutaneous injection into the right flank, proximal to the hip, with 100*μ*l of an emulsion of Ovalbumin (OVA) with incomplete Freunds adjuvant (IFA) (100*μ*g OVA per mouse) (Hooke Laboratories). Where stated, mice were treated with Y-27632 (Tocris 1254). 10*μ*l of 1mg/ml Y-27632 dissolved in PBS was injected subcutaneously into the right flank, proximal to the hip. 10*μ*l of sterile PBS was used as injection control. Y-27632/PBS injections were given on 3 consecutive days 24 hours post IFA/OVA injection. After 5 or 3 days mice were culled and inguinal lymph nodes (LNs) from both flanks (naïve and inflamed) were extracted for paired histological studies, flow cytometry analysis or ex vivo laser ablation.

#### Ex vivo cultures

Preparation for ex vivo lymph nodes laser ablation was optimised following previously established methods UltraPure™ Low melting point agarose (Thermo Fischer Scientific) prepared in PBS at 3% w/v and maintained at 37°C. Lymph nodes were embedded into agarose and left for 5 minutes to set on ice. Lymph node blocks were secured by superglue to cutting stage and placed into ice cold PBS cutting chamber. Lecia Vibratome (VT1200S) 200*μ*m thick sections were cut at a rate of 0.3mm/s and 1.5mm amplitude until the LN was completely sectioned. Collected sections were placed into RPMI 1640 containing 10% FBS, 1% penicillin and streptomycin (P/S) (Thermo Fischer scientific 15140122) and 1% Insulin-Transferrin-Selenium (ITS) (Thermo Fischer scientific 41400045) at 37°C, 10% CO_2_. Sections recovered for an hour before been used for live imaging.

#### Laser ablation

LN sections were transferred to glass bottom 35mm MatTek dishes (P35G-1.5-20-C) with a small volume of RPMI media containing 1% P/S and 1% ITS. A glass cover slip was placed on top to secure the section. Sections imaged on a Zeiss LSM 880 inverted multiphoton microscope with the imaging chamber maintained at 37°C, 10% CO_2_. Where stated, sections were treated with 100 *μ*M Y-27632 (Tocris 1254) diluted in *ex vivo* culture media for at least 1 hour prior to imaging and ablation. Sections were imaged with a Plan Apochromat 40x oil objective (NA1.3), 1024*1024 resolution and 4x digital zoom for ablation ROIs. Laser ablation of the FRC network was achieved by using a pulsed Chameleon Vision II TiSa Laser power 75-80% (coherent) tuned to 760nm. Ablation was performed on small, manually defined linear regions of interest between FRC connections away from cell bodies. Ablation was performed in a single z-plane at the centre of the FRC connection in a vibratome slice. Time-lapse videos were recorded over 25 seconds on a single channel (PMT detector) 512*512 pixels, with 521 millisecond scan speed per frame to capture recoil of the network. Recoil of the network was calculated by manually measuring displacement of the FRC network between two points located away from the ablation site, with initial recoil velocities calculated from the displacement one frame after the cut, as in other studies^7,8^.

#### Immunostaining of tissue sections

LNs that were used for sectioning and immunofluorescence were fixed in Antigen fix (DiaPath) for 2 hours at 4°C with gentle agitation. LNs were washed for 30 minutes in PBS before being applied to 30% w/v sucrose 0.05% sodium azide solution at 4°C overnight. LNs were dipped into Tissue-Tek OCT before being embedded into moulds containing OCT. A maximum of six lymph nodes were embedded into a single block for comparative analysis. Lymph nodes were sectioned on the Leica cryostat at a thickness of 20um.

For immunofluorescent staining sections were permeabilised and blocked using 10% Normal goat serum (NGS), 0.3% Triton X-100 in PBS for two hours at room temperature. Primary antibodies were prepared by centrifugation at 13,000rpm for 5 minutes at 4°C. Primary antibodies were diluted according to ***Antibody Table 1*** in 10% NGS, 0.01% Triton X-100 in PBS. Sections were incubated overnight at 4°C. Sections were then brought to room temperature and washed 3 times for 15 minutes each in 0.05% PBS-Tween20. Sections were then blocked using 10% Normal goat serum (NGS), 0.3% Triton X-100 in PBS for two hours at room temperature. Secondary antibodies were then prepared similar to primary antibodies with dilutions in ***Antibody Table 1*.** Secondary antibodies were applied to sections for 2hrs at room temperature. This was followed by two 15-minute washes of 0.05% PBS-Tween20 and a final wash of 15 minutes in PBS. Sections were then mounted using mowiol mounting media.

For staining of 200*μ*m thick vibratome sections, slices were first fixed in Antigen fix (DiaPath) for 2 hours at 4°C with gentle agitation. Sections were placed into 0.1M Tris-HCL pH7.4 on ice for 30 minutes. Sections were permeabilised using IHC buffer containing 0.5% BSA, 2% Triton X-100 (Sigma-Aldrich) in 0.1M Tris-HCL pH7.4 for 20 minutes at 4°C with gentle agitation. Primary antibodies were prepared by centrifugation at 13,000rpm for 5 minutes at 4°C. Primary antibodies were diluted into IHC buffer, according to ***Antibody Table 1*,** and applied to the sections for 1hr at 4°C with gentle agitation. Sections were then washed twice with 0.1M Tris-HCL for 15 minutes. Secondary antibodies were then prepared similar to primary antibodies with dilutions in ***Antibody Table 1*.** Secondary antibodies were applied to sections for 1hr at 4°C with gentle agitation. Two final washes of 0.1M Tris-HCL were performed before mounting the sections with mowiol mounting media and a glass coverslip. Sections (15-3=40 *μ*m) were then imaged on the Zeiss LSM 880 inverted multiphoton microscope using a Plan Apochromat 40x oil objective (NA1.3).

Unless otherwise stated confocal images were acquired using Leica TCS SP8 STED 3X or the Leica TCS SP5. Images were captured at 1024*1024 pixels, 3 line average onto HyD or PMT detectors. Fluorophore excitation and acquisition was performed in a sequential and bidirectional manner. Imaging regions were manually defined and z-stacks (15-40 *μ*m) with regular z-intervals ranging from 0.4um or 1um (dependent on sample) were acquired using a motorised stage. Tile scans were automatically stitched (numerical) using Leica imaging software.

#### Flow cytometry of lymph nodes

LNs were carefully dissected, weighed, and placed into RPMI 1640 media on ice. LNs were then processed as previously described ^9,10^. Briefly, LNs were placed into a digestion buffer containing collagenase D (250*μ*g/ml) (Millipore Sigma), dispase II (800*μ*g/ml) (Thermo Fisher Scientific) and DNase I (100*μ*g/ml) (Sigma Aldrich). LNs were gently digested in a water bath at 37 °C, removing and replacing the cell suspension every 10 minutes until completely digested. Cell suspensions were then centrifuged at 350g for 5 minutes. The cells were resuspended into flow cytometry buffer (FACS) consisting of 1% BSA (Sigma Aldrich), 5mM EDTA (Sigma Aldrich) in PBS, filtered, counted, and resuspended at 10 × 10^6^ *cells/ml*. 2.5 × 10^6^ cells were then plated into 96 well plates for surface and intracellular staining of a stromal cell panel **(*Antibody Table 1*).** Cells were blocked with CD16/CD32 Mouse Fc block (BD) and then stained with primary antibodies for 20 minutes at 4°C. For intracellular staining of Ki67 cells were fixed and permeabilised using FOXP3 fix/perm buffer as specified by the manufacturer (BioLegend). Samples were run on the Fortessa X20 flow cytometer (BD Biosciences) at the UCL Cancer Institute. Data was analysed using Flow Jo software (FlowJo, LLC).

#### Cell lines

Immortalised fibroblastic reticular cells were generated as described in Acton et al 2014^10,11^. Parental immortalised fibroblastic reticular cell line (Control FRC). Podoplanin (PDPN) was stably knocked down (PDPN KD FRC) in the parental cell line by transfection of a PDPN shRNA lentivirus. PDPN was completely depleted from the parental cell line (PDPN KO FRC) using CRISPR cas9 genetic deletion. In all experiment where exogenous PDPN mutant cell lines (PDPN WT, PDPN T34A, PDPN S167A-S171A) were used, protein production was induced by the addition of 1*μ*g/ml of doxycycline for 48 hours. Cell lines were maintained at 10% CO_2_, 37°C in Dulbecco’s modified eagle medium (DMEM) (Thermo Fischer scientific) supplemented with 10% FBS, 1% P/S and 1% ITS unless otherwise stated. Cells were treated with recombinant CLEC2-Fc or Control-Fc supernatant ^10,11^ for approximately 2hrs, where indicated, to assay the effect of CLEC-2 signalling through PDPN. Figure 3C is a paired analysis of the same cell before and 15 minutes after CLEC-2 treatment.

#### Tether pulling and trap force measurements

Trap force measurements were performed using a home built optical tweezer using a 4W 1064 nm laser quantum Ventus within a 100x oil immersion objective (NA 1.30, CFI Plan Fluor DLL, Nikon) on an inverted microscope (Nikon Eclipse TE2000-U) equipped with a motorised stage (PRIOR Proscan). The optical tweezer was calibrated following previous studies ^12,13^. The trap force calibration was performed in every experiment with typical calibration trap stiffness of k~0.114 *pN/nm*. Measurements were performed using concanavalin-A coated (50ug/ml) carboxyl latex beads, (1.9um diameter, Thermo Fischer). Beads were incubated on a shaker with concanavalin-A for two hours prior to experiment. Beads were applied to the culture media, manipulated by the optical trap and brought into contact with the cellular membrane and typically held for 2-5 seconds to allow binding to membrane. Bead position was recorded every 90 milliseconds in bright field prior and during tether formation. Cells were maintained in the trap at 37 degrees and had CO_2_ flowing into the chamber. Addition of CLEC-2 or CTRL supernatant was 2-4 hours prior to measurement of trap force. Trap force (F_t_ - *pN/*μ*m*) was then calculated based on the trap calibration (*k*), bead position (Δ*x*) using a home-made Fiji macro ^13^ and the equation *F_T_* = *k*Δ*x*.

#### Omsotic swelling assay

Osmotic shock was performed in accordance with previously reported protocols ^14^. By altering the osmolarity of a solution cells will swell or shrink. Osmolarity was estimated using osmolarity calculations. Isotonic solution was prepared with 137mM NaCl, 5.4mM KCl, 1.8mM CaCl_2_, 0.8mM MgCl_2_, 20mM HEPES, 20mM D-glucose and pH to 7.4 with NaOH. Hypotonic (Hypo) solutions were prepared by diluting the isotonic preparation in milliq water i.e. 50 mOsm is a 1/6 dilution of Iso. To control for the dilution of ions in solution and account for the effect this may have on swelling, the 50 mOsm solution was restored to 330mOsm using D-mannitol at 280mM, acting as the true isotonic control (Iso). Cells were dissociated from cell culture with Sterile Dulbecco’s phosphate buffered saline (DPBS) (Thermo Fischer scientific) + 2mM EDTA and placed onto individual 35mm MatTek dishes and allowed to settle for 30 minutes. After 30 minutes the cells remain rounded on the coverslip. Cells were then treated with either Iso, Hypo 50mOsm or Extreme Hypo 0mOsm for one hour. Phase contrast images of cell swelling were captured every 30 seconds using 20x air objective on a Nikon Ti inverted microscope with a motorised stage controlled by NIS-elements software. Diameter of individual swelling cells were calculated using manual circular ROIs. The ratio of swelling was then calculated by dividing all diameters (d) by the initial diameter (d0). Area under the curves were calculated for the first 20 minutes of the swelling response.

#### Western blotting

Equal cell numbers were grown to confluency and protein was isolated using 300*μ*l of 4x Laemmlli Lysis Buffer (Bio-Rad) and cell lifters (Fisher Scientific). Lysates were then sonicated for 20 seconds followed by 10 minutes at 95°C. 1% β-mercaptoethanol (143mM stock, Sigma Aldrich) was applied to samples to reduce oligomerised protein structures. Electrophoresis gels were loaded with the same quantity of lysates and run for 60 minutes at 110v. Transfer to PDVF membranes were carried out at 65v for 2hr at 4. Membranes were blocked for 2hrs at room temperature with 5% skim milk powder (Sigma-Aldrich), 1% BSA in TBS and stained with primary antibodies **(*Antibody Table 1*)** overnight at 4 in 1:5 diluted blocking buffer. Membranes were then washed thoroughly with TBS 0.05% Tween 20 and incubated with HRP-conjugated secondary antibodies (Antibody Table 1) in 1:5 blocking buffer. After washing with TBS 0.05% Tween 20. The membranes were visualised using ECL-HRP reaction and imaged using Image Quant 5000 (GE Lifesceinces).

#### Linear unmixing and Imaris rendering

Imaging of PDGFR*α*-mGFP CreERT2 Confetti LN sections was carried out using lambda mode and chameleon laser at 900um to acquire multi-channel images. Widefield images and z-stack intervals of 0.5mm were obtained, for an approximate thickness of 30-60uM. The emission wavelengths for each fluorophore were set on Zeiss zen black software spectral unmixing function by selecting labelled cells within the confetti imaging. The 2nd harmonic of the two-photon laser detected the ECM conduit structure in the lymph node. Rendering of PDPN, CFP and YFP in Fig. 1, 2, 5, Fig. S5 was achieved using Imaris surface tools.

#### Quantification and statistical analysis

Prism7 Software (GraphPad) was used to perform multiple statistical analyses including appropriate tests were performed as indicated in figure legends. In general, comparison of multiple groups was performed using Two-way ANOVA with Tukey’s multiple comparisons or One-way ANOVA with Kruskal-Wallis test depending on the distribution of the data. Comparisons of two data sets were most performed using Two-tailed Mann-Whitney tests.

**Fig. S1 -.**
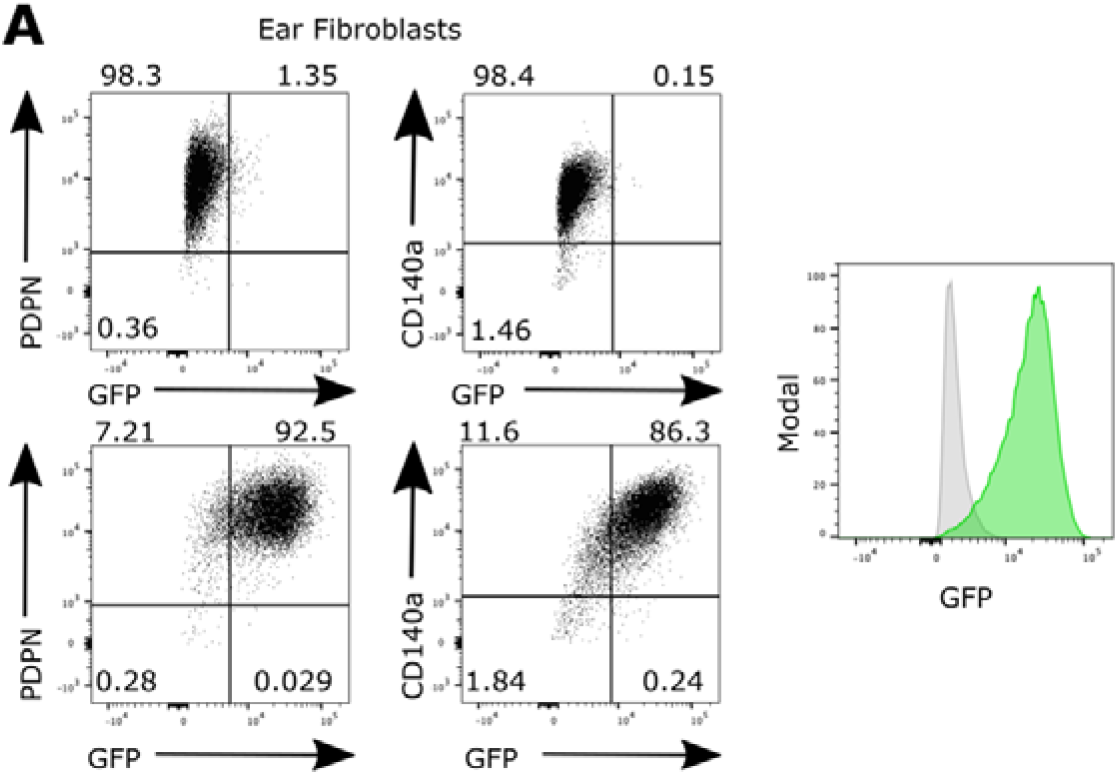
Flow cytometry of PDGFR -mGFP fibroblasts. **A)** Flow cytometry panel of PDPN^+^ and CD140 ^+^ skin fibroblasts expressing GFP. Top two panels show GFP-control fibroblasts. Bottom two panels show PDPN+, GFP+ and CD140a+ GFP+ fibroblasts. Right panel shows control fibroblasts (grey) to PDGFR -mGFP fibroblasts (green).

**Fig. S2 –.**
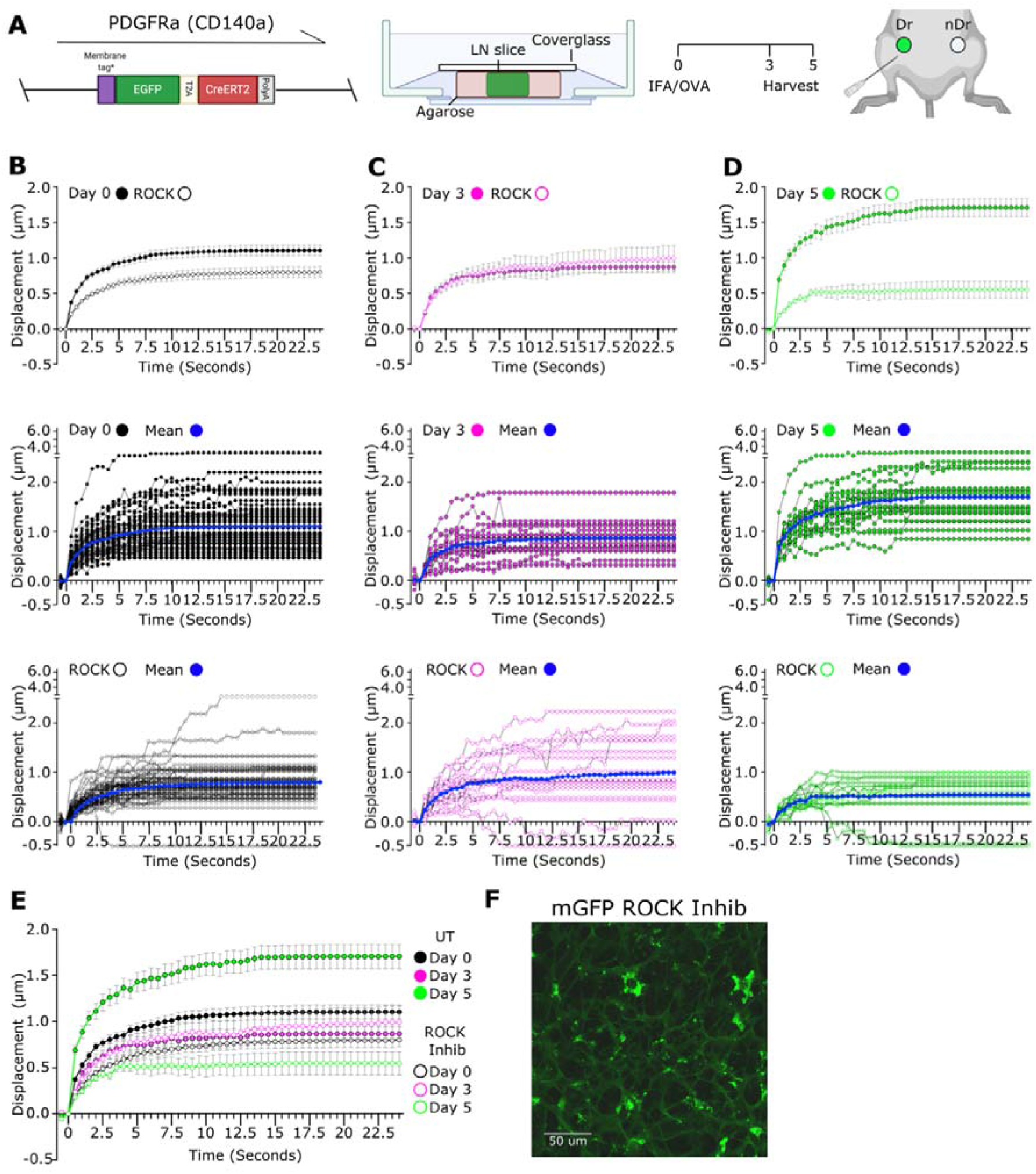
Y27 treatment of lymph node slices reduce tension in the FRC network. **A)** Schematic of immunisation and animal models. A membrane targeted GFP molecule is driven under the (PDGFR) promoter (left). LN nodes were embedded in low melt agarose and sliced at 200 m thickness before being secured by cover glass for imaging (middle). IFA/OVA is used as model immunisation with inguinal draining (dr) and non-draining (nDr) lymph nodes (LN’s) harvested day 3 or day 5 post immunisation (right). **B-D)** Recoil curves show displacement over time (mean ±SEM) (top panel). Individual recoil curves of control (middle) and Y27 treated LNs (bottom). N>5 animals per condition. **E)** Recoil curves show displacement over time (mean ±SEM), comparing untreated and Y27 treated LNs. Blue shading indicates initial recoil phase. N>5 animals per condition. **F)** Membrane GFP visualises the FRC network in the *ex vivo* lymph node slice. Y27 treatment has no effect on the FRC network connectivity. Scale bar 50 m.

**Fig. S3 -.**
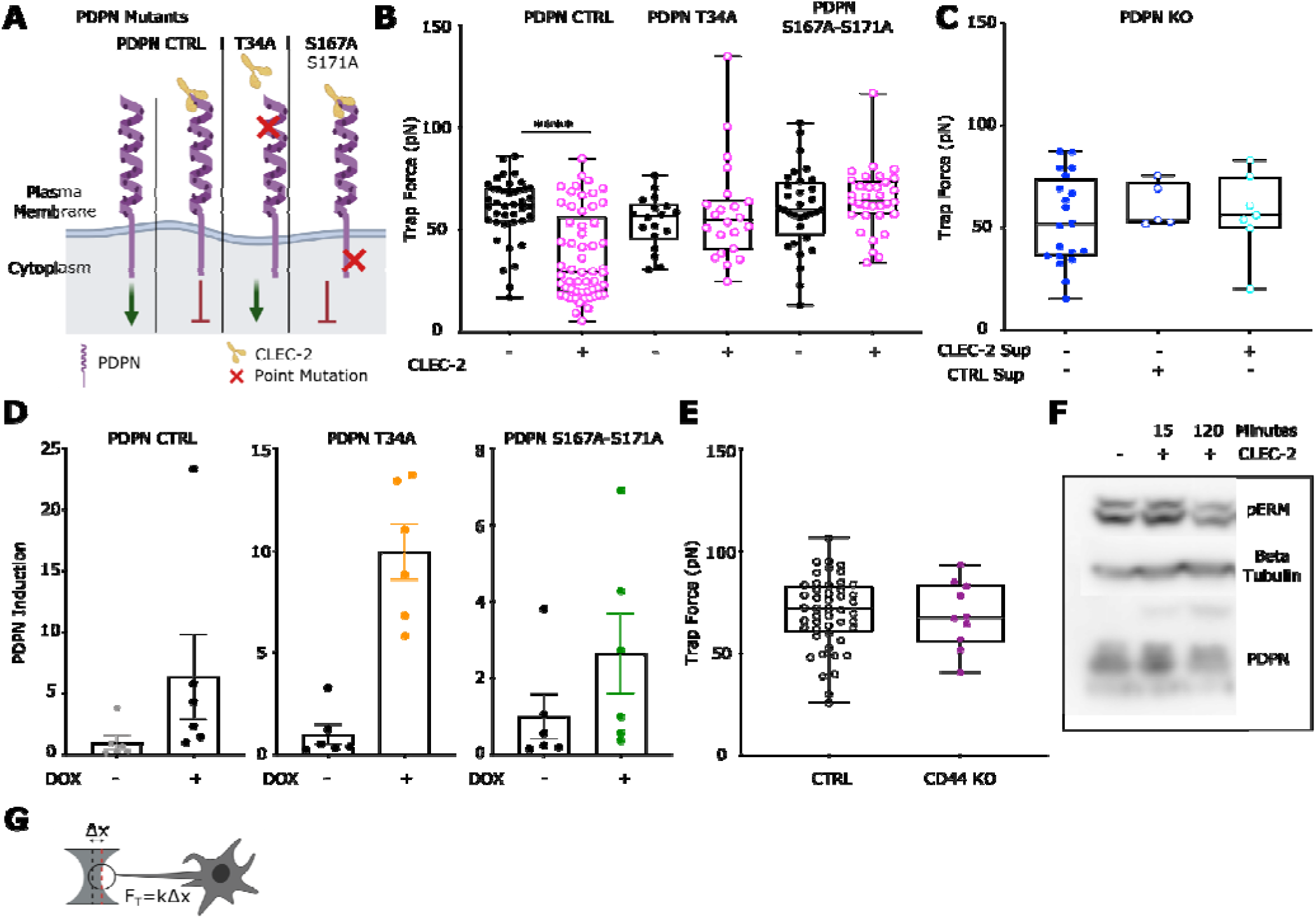
Induction of PDPN mutants in PDPN KO fibroblastic reticular cell line. **A)** Schema of exogenous PDPN mutants and interaction with CLEC2. Green arrow denotes active signalling by PDPN leading to actomyosin contractility, red arrow indicates inhibition of PDPN signalling and reduction in actomyosin contractility. **B)** Trap force measurements of FRCs expressing PDPN mutants after pre-treatment of CLEC2. Box plots indicates median and interquartile range. Two-way ANOVA with Tukeys multiple comparisons, p<0.001. N>18. **C)** Trap force measurements of PDPN CRISPR KO FRCs treated with CTRL or CLEC2 supernatant. One-way ANOVA with Tukeys multiple comparisons. N>5. **D)** Fold induction of exogenous PDPN, based on PDPN staining geometric mean, for each PDPN mutant cell line treated with or without doxycycline. PDPN CTRL (left), PDPN T34A (middle) and PDPN DSS (right). N=6. **E)** Trap force measurements of CTRL and CD44 (purple) KO FRCs. Mann-Whitney test, p<0.001. Each point represents one cell. N>10. **F)** Representative western blot of pERM in CTRL FRCs after treatment with CLEC2. **G)** Schema and equation to calculate trap force (F_t_) from displacement.

**Fig. S4 -.**
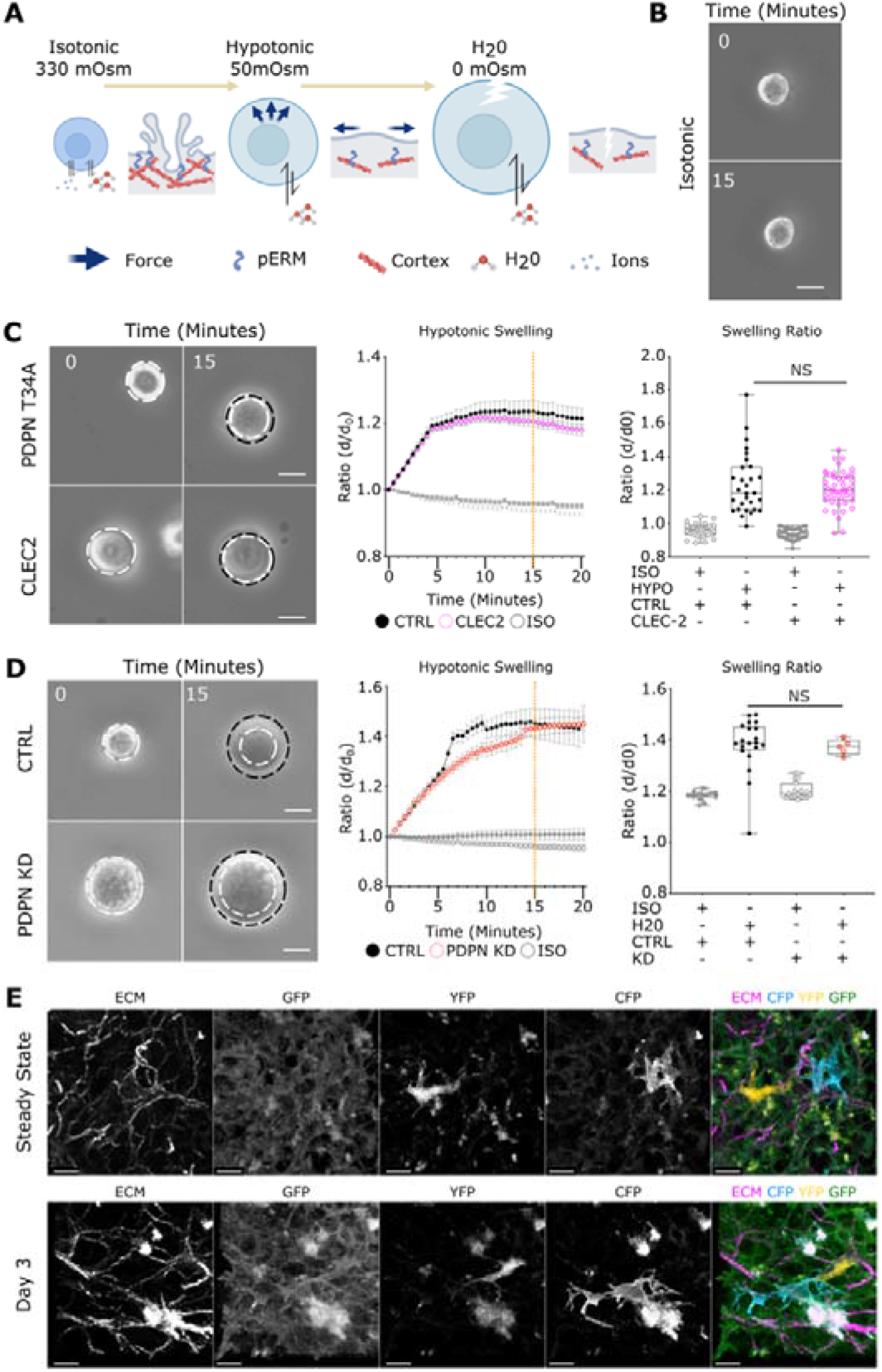
Membrane access of fibroblastic reticular cells is increased through the CLEC2/PDPN signalling axis. **A)**Schematic of osmotic shock experiment, inducing swelling by altering osmolarity. **B)** Representative stills of FRC in isotonic control media. Scale bar 25 m. **C)** Time course of PDPN T34A FRCs with or without CLEC2 treatment in hypotonic conditions. Stills (top) show swelling of cells in hypotonic media. White dotted circle marks initial size before swelling and is compared to swelling at t=20 (black dotted circle). Scale bar 25 m. Change in diameter ratio over time (bottom left, mean ±SEM). Swelling ratio comparisons between control and CLEC2 treatment at 15 minutes post swelling (orange dotted line, bottom right). Box plots indicates median and interquartile range. One-way ANOVA with Tukeys multiple comparisons. N>29. **D)** Time course of PDPN CTRL vs PDPN shRNA KD FRCs in extreme hypotonic conditions (H_2_0). Stills (top) show swelling of cells in hypotonic media. White dotted circle marks initial size before swelling and is compared to swelling at t=20 (black dotted circle). Scale bar 25 m. Change in diameter ratio over time (bottom left, mean ±SEM). Swelling ratio between control and PDPN shRNA KD FRCs at 15 minutes post swelling (orange dotted line, bottom right). Box plots indicates median and interquartile range. One-way ANOVA with Tukeys multiple comparisons. N>16. **E)** Single channel stills of PDGFR -mGFP confetti FRC network in the steady state and day 3 post immunisation. Scale bar 10 m.

**Fig. S5 -.**
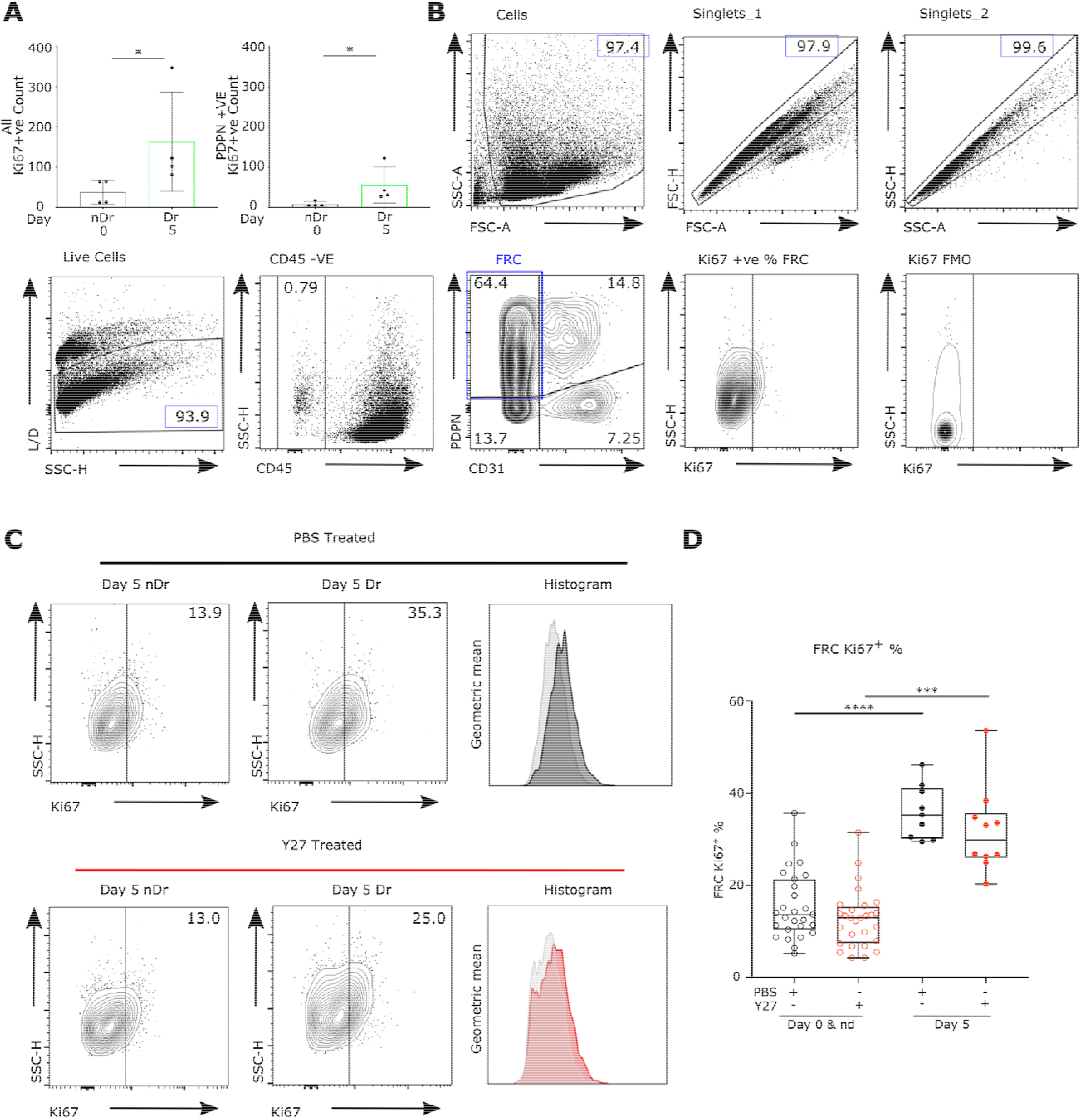
Flow cytometry gating strategy and Ki67 measurements. **A)** Quantification of Ki67^+^ cells (top) and PDPN^+^ Ki67^+^ cells (bottom) per LN region. Box plots indicates median and interquartile range. Mann-Whitney test, p<0.05. N=2. **B)** Schema of flow cytometry gating strategy for measurement of stromal cell populations. **C)** Example of Ki67^+^ FRCs cell plots and histogram of Ki67 geometric mean, comparing immunisation and Y27 treatments. **D)** Quantification of Ki67^+^ FRCs. Box plots indicates median and interquartile range. Two-way ANOVA with Tukey’s multiple comparisons. ****p<0.0001, ***p<0.001. N>9 animals per condition

**Table S1 –.**
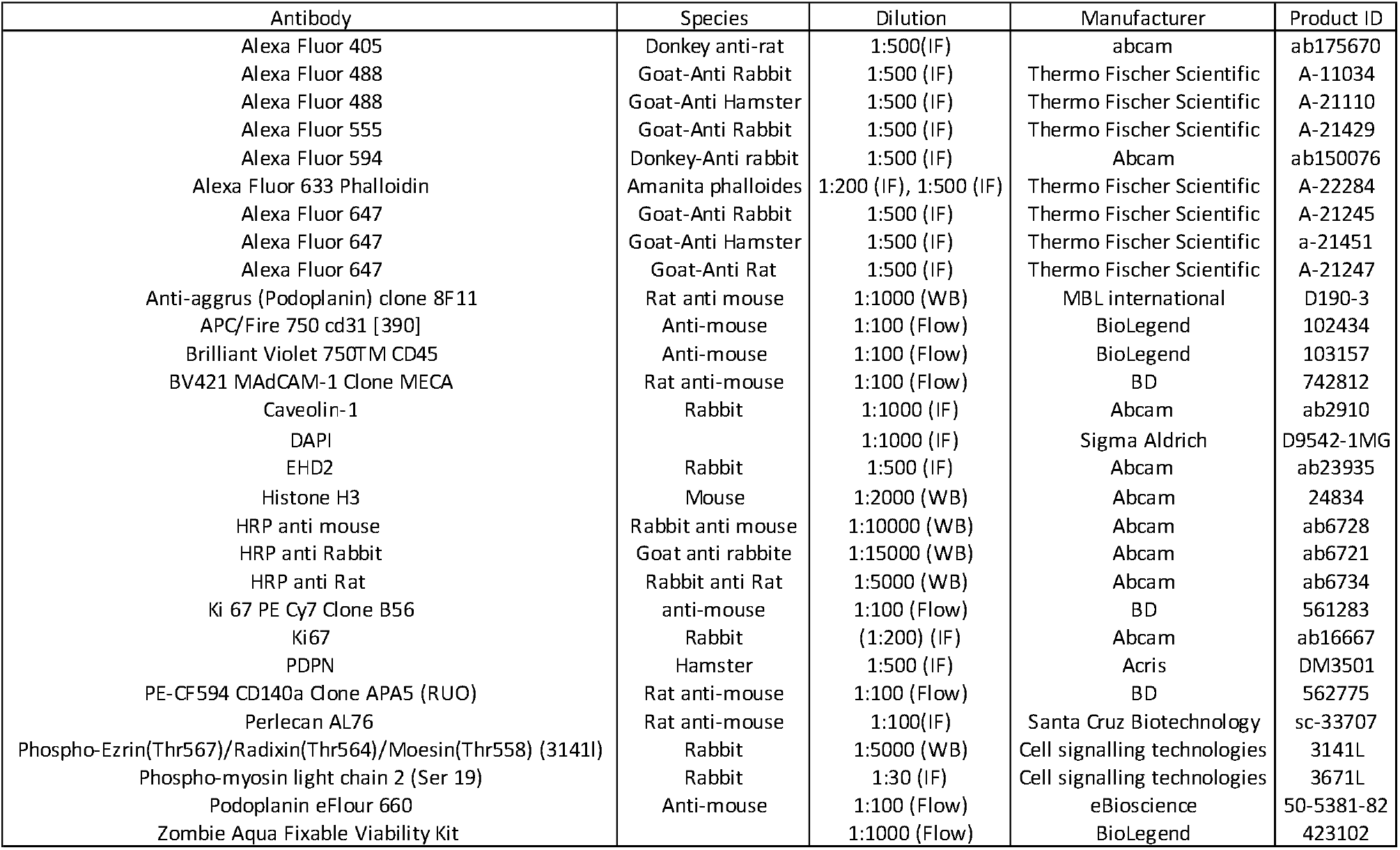
Antibody reagents and dilutions.

| Antibody | Species | Dilution | Manufacturer | Product ID |
| --- | --- | --- | --- | --- |
| Alexa Fluor 405 | Donkey anti-rat | 1:500(IF) | abcam | ab175670 |
| Alexa Fluor 488 | Goat-Anti Rabbit | 1:500 (IF) | Thermo Fischer Scientific | A- 11034 |
| Alexa Fluor 488 | Goat-Anti Hamster | 1:500 (IF) | Thermo Fischer Scientific | A- 21110 |
| Alexa Fluor 555 | Goat-Anti Rabbit | 1:500 (IF) | Thermo Fischer Scientific | A- 21429 |
| Alexa Fluor 594 | Donkey-Anti rabbit | 1:500 (IF) | Abcam | ab150076 |
| Alexa Fluor 633 Phalloidin | Amanita phalloides | 1:200 (IF), 1:500 (IF) | Thermo Fischer Scientific | A- 22284 |
| Alexa Fluor 647 | Goat-Anti Rabbit | 1:500 (IF) | Thermo Fischer Scientific | A- 21245 |
| Alexa Fluor 647 | Goat-Anti Hamster | 1:500 (IF) | Thermo Fischer Scientific | a- 21451 |
| Alexa Fluor 647 | Goat-Anti Rat | 1:500 (IF) | Thermo Fischer Scientific | A- 21247 |
| Anti-aggrus (Podoplanin) clone 8F11 | Rat anti mouse | 1:1000 (WB) | MBL international | D190-3 |
| APC/Fire 750 cd31 [390] | Anti-mouse | 1:100 (Flow) | BioLegend | 102434 |
| Brilliant Violet 750TM CD45 | Anti-mouse | 1:100 (Flow) | BioLegend | 103157 |
| BV421 MAdCAM-1 Clone MECA | Rat anti-mouse | 1:100 (Flow) | BD | 742812 |
| Caveolin-1 | Rabbit | 1:1000 (IF) | Abcam | ab2910 |
| DAPI |  | 1:1000 (IF) | Sigma Aldrich | D9542-1MG |
| EHD2 | Rabbit | 1:500 (IF) | Abcam | ab23935 |
| Histone H3 | Mouse | 1:2000 (WB) | Abcam | 24834 |
| HRP anti mouse | Rabbit anti mouse | 1:10000 (WB) | Abcam | ab6728 |
| HRP anti Rabbit | Goat anti rabbite | 1:15000 (WB) | Abcam | ab6721 |
| HRP anti Rat | Rabbit anti Rat | 1:5000 (WB) | Abcam | ab6734 |
| Ki 67 PE Cy7 Clone B56 | anti-mouse | 1:100 (Flow) | BD | 561283 |
| Ki67 | Rabbit | (1:200) (IF) | Abcam | ab16667 |
| PDPN | Hamster | 1:500 (IF) | Acris | DM3501 |
| PE-CF594 CD140a Clone APA5 (RUO) | Rat anti-mouse | 1:100 (Flow) | BD | 562775 |
| Perlecan AL76 | Rat anti-mouse | 1:100(IF) | Santa Cruz Biotechnology | sc-33707 |
| Phospho-Ezrin(Thr567)/Radixin(Thr564)/Moesin(Thr558) (3141l) | Rabbit | 1:5000 (WB) | Cell signalling technologies | 3141L |
| Phospho-myosin light chain 2 (Ser 19) | Rabbit | 1:30 (IF) | Cell signalling technologies | 3671L |
| Podoplanin eFlour 660 | Anti-mouse | 1:100 (Flow) | eBioscience | 50-5381-82 |
| Zombie Aqua Fixable Viability Kit |  | 1:1000 (Flow) | BioLegend | 423102 |

## Movies S1 to S4

**S1A – The paracortical T-cell FRC network, showing 3D localisation of actomyosin structures in homeostasis.**

Staining of the conduit (perlecan, magenta), FRC network (PDPN, yellow), F-actin (phalloidin, cyan) and pMLC (red). PDPN and perlecan surface renders show the location of actomyosin in relation to these structures.

**S1B – Laser ablation of homeostatic FRC network.**

Movies captured over 25 seconds. Yellow line marks site of cut one second after imaging began.

**S2A – Laser ablation of FRC network day 3 post immunisations.**

Movies captured over 25 seconds. Yellow line marks site of cut one second after imaging began.

**S2B – Laser ablation of FRC network day 5 post immunisations.**

Movies captured over 25 seconds. Yellow line marks site of cut one second after imaging began.

**S2C – The paracortical T-cell FRC network, showing 3D localisation of actomyosin structures day 3 post immunisations.**

Staining of the conduit (perlecan, magenta), FRC network (PDPN, yellow), F-actin (phalloidin, cyan) and pMLC (red). PDPN and perlecan surface renders show the location of actomyosin in relation to these structures.

**S2D – The paracortical T-cell FRC network, showing 3D localisation of actomyosin structures day 5 post immunisation.**

Staining of the conduit (perlecan, magenta), FRC network (PDPN, yellow), F-actin (phalloidin, cyan) and pMLC (red). PDPN and perlecan surface renders show the location of actomyosin in relation to these structures.

**S3A – Optical tweezers generate membrane tethers used to determine membrane tension**

**S4A – Osmotic swelling of PDPN CTRL fibroblastic reticular cells**

PDPN CTRL + CLEC2 (left), PDPN CTRL – CLEC2 (middle), PDPN CTRL ISO control (right). Captured over 60 minutes with one frame every 30 seconds.

**S4B – Osmotic swelling of PDPN CTRL fibroblastic reticular cells**

B) PDPN T34A + CLEC2 (left), PDPN T34A – CLEC2 (middle), PDPN T34A ISO control (right). Captured over 60 minutes with one frame every 30 seconds.

**S4C – Osmotic swelling of PDPN CTRL fibroblastic reticular cells**

**C)** CTRL + HYPO (left), PDPN KD + HYPO (middle), CTRL ISO control (right). Captured over 60 minutes with one frame every 30 seconds.

**S4D – Osmotic swelling of PDPN CTRL fibroblastic reticular cells**

**D)** CTRL + H_2_0 (left), PDPN KD + H_2_0 (right). Captured over 60 minutes with one frame every 30 seconds.

