## Supplementary figures for "Tissue homeostasis and adaptation to immune challenge resolved by fibroblast network mechanics"

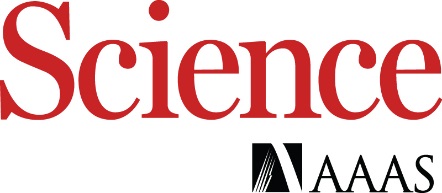

Supplementary Materials for

**Tissue homeostasis and adaptation to immune challenge resolved by fibroblast network mechanics**

Harry L. Horsnell^1^, Robert J. Tetley^2^, Henry De Belly^3^, Spyridon Makris^1^, Lindsey J. Millward^1^, Agnesska C. Benjamin^1^, Charlotte M de Winde^1^, Ewa K. Paluch^3^, Yanlan Mao^2,4^, Sophie E. Acton^1#^

**This PDF file includes:**

Materials and Methods

Figs. S1 to S5

Tables S1

References (1 to 14)

Captions for Movies S1 to S4

**Other Supplementary Materials for this manuscript include the following:**

**Figs. S1 to S5**

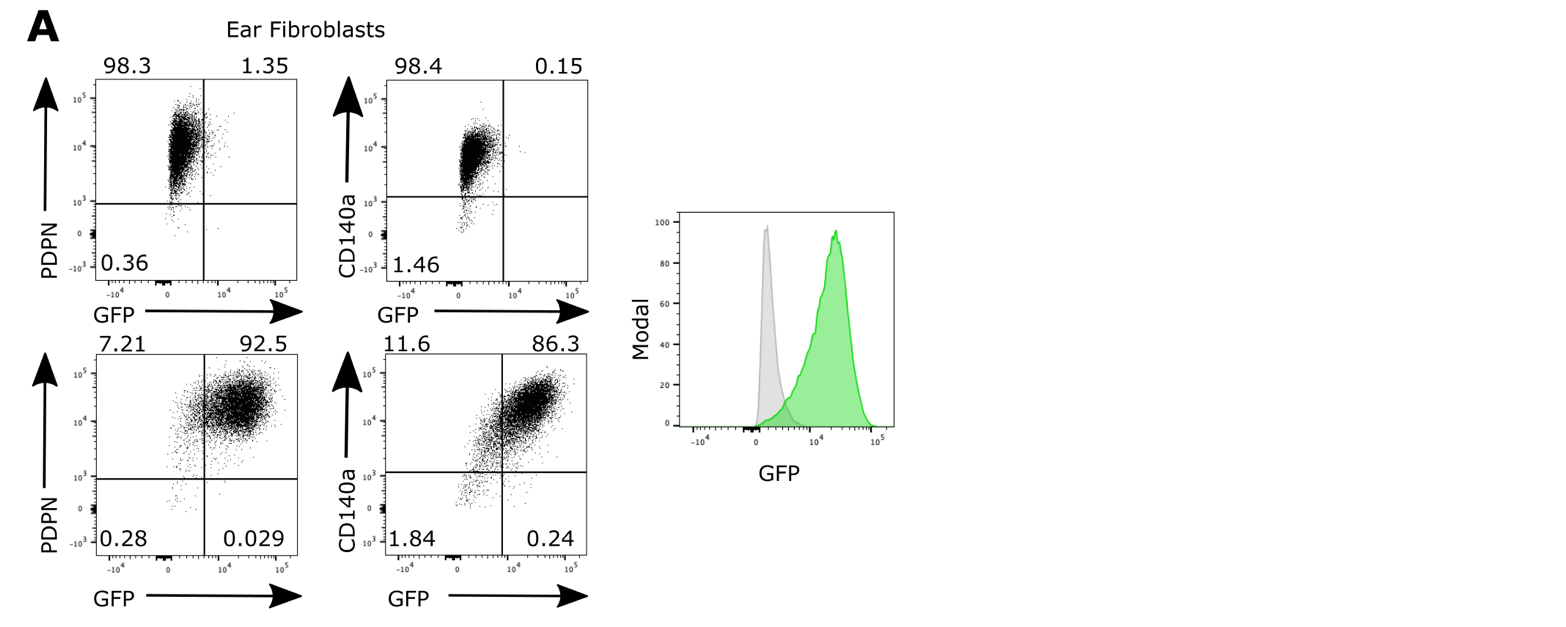

**Fig. S1** - **Flow cytometry of** **PDGFR**$\boldsymbol{\alpha}$**-mGFP fibroblasts**

**A)** Flow cytometry panel of PDPN^+^ and CD140$\alpha$^+^ skin fibroblasts expressing GFP. Top two panels show GFP- control fibroblasts. Bottom two panels show PDPN+, GFP+ and CD140a+ GFP+ fibroblasts. Right panel shows control fibroblasts (grey) to PDGFR$\alpha$-mGFP fibroblasts (green).

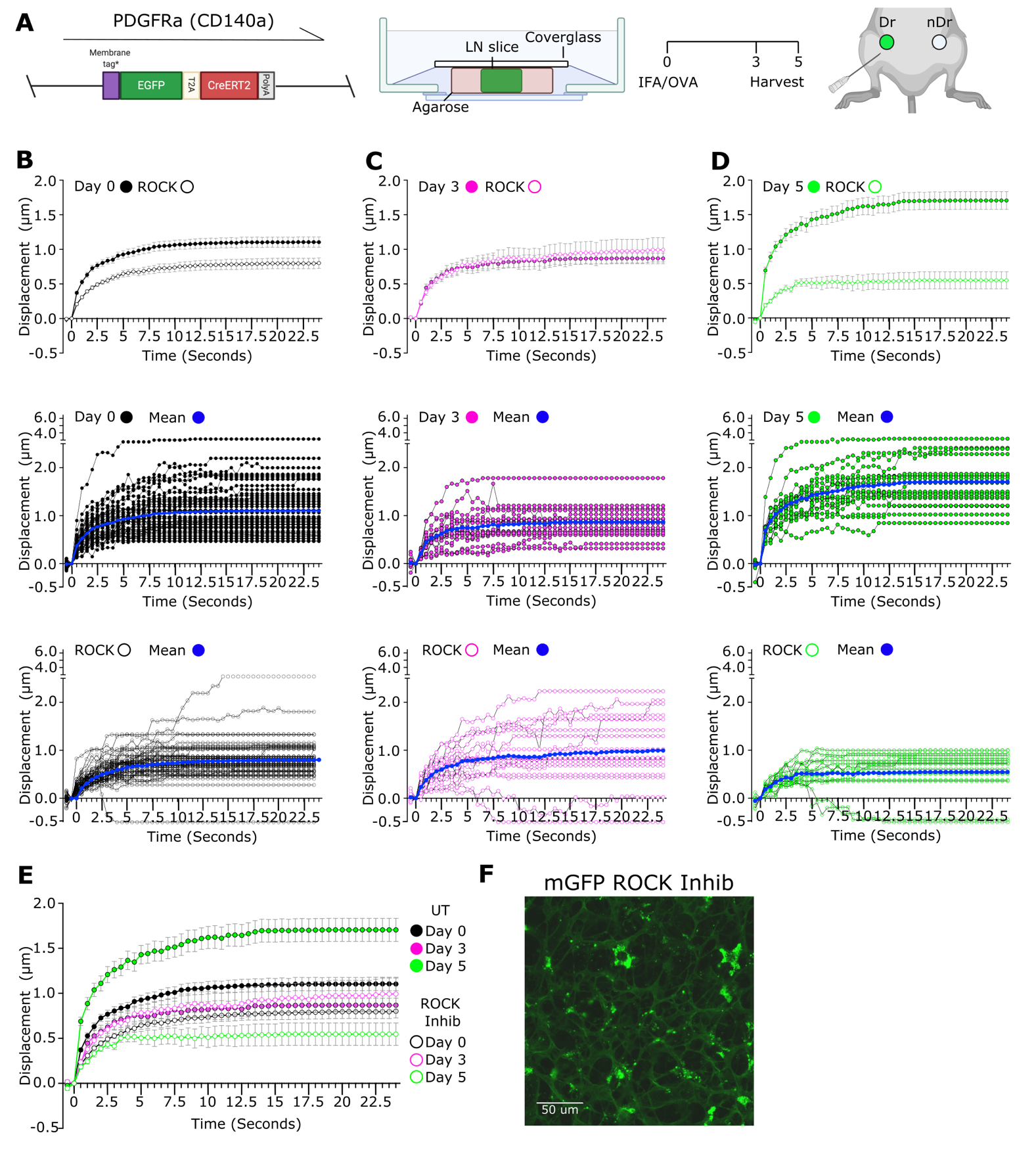

**Fig. S2** – **Y27 treatment of lymph node slices reduce tension in the FRC network**

**A)** Schematic of immunisation and animal models. A membrane targeted GFP molecule is driven under the (PDGFR$\alpha$) promoter (left). LN nodes were embedded in low melt agarose and sliced at 200$\mu$m thickness before being secured by cover glass for imaging (middle). IFA/OVA is used as model immunisation with inguinal draining (dr) and non-draining (nDr) lymph nodes (LN’s) harvested day 3 or day 5 post immunisation (right). **B-D)** Recoil curves show displacement over time (mean ±SEM) (top panel). Individual recoil curves of control (middle) and Y27 treated LNs (bottom). N>5 animals per condition. **E)** Recoil curves show displacement over time (mean ±SEM), comparing untreated and Y27 treated LNs. Blue shading indicates initial recoil phase. N>5 animals per condition. **F)** Membrane GFP visualises the FRC network in the *ex vivo* lymph node slice. Y27 treatment has no effect on the FRC network connectivity. Scale bar 50$\mu$m.

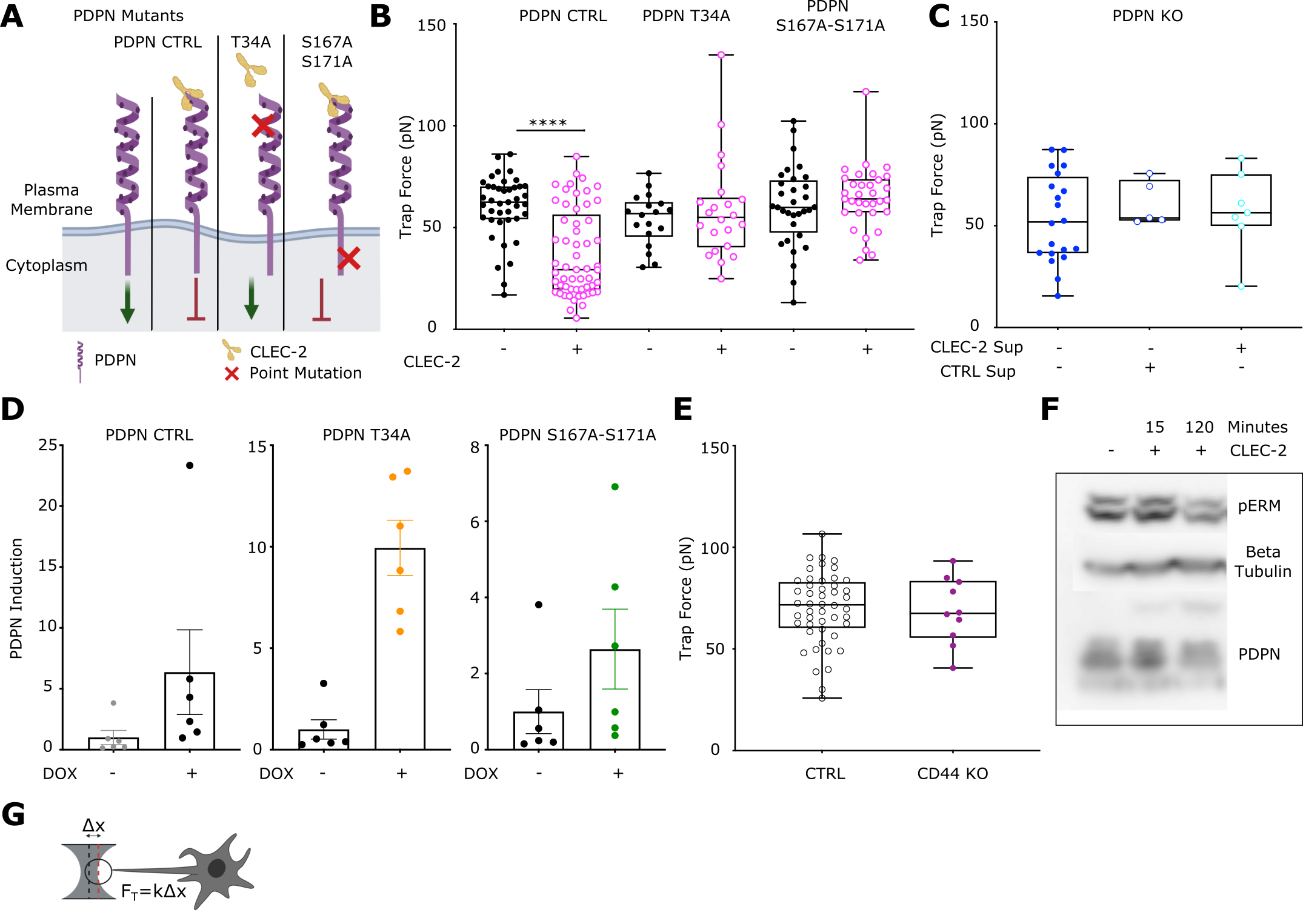

**Fig. S3 - Induction of PDPN mutants in PDPN KO fibroblastic reticular cell line**

**A)** Schema of exogenous PDPN mutants and interaction with CLEC2. Green arrow denotes active signalling by PDPN leading to actomyosin contractility, red arrow indicates inhibition of PDPN signalling and reduction in actomyosin contractility**. B)** Trap force measurements of FRCs expressing PDPN mutants after pre-treatment of CLEC2. Box plots indicates median and interquartile range. Two-way ANOVA with Tukeys multiple comparisons, p<0.001. N>18**. C)** Trap force measurements of PDPN CRISPR KO FRCs treated with CTRL or CLEC2 supernatant. One-way ANOVA with Tukeys multiple comparisons. N>5**. D)** Fold induction of exogenous PDPN, based on PDPN staining geometric mean, for each PDPN mutant cell line treated with or without doxycycline. PDPN CTRL (left), PDPN T34A (middle) and PDPN DSS (right). N=6. **E)** Trap force measurements of CTRL and CD44 (purple) KO FRCs. Mann-Whitney test, p<0.001. Each point represents one cell. N>10**. F)** Representative western blot of pERM in CTRL FRCs after treatment with CLEC2**. G)** Schema and equation to calculate trap force (F_t_) from displacement.

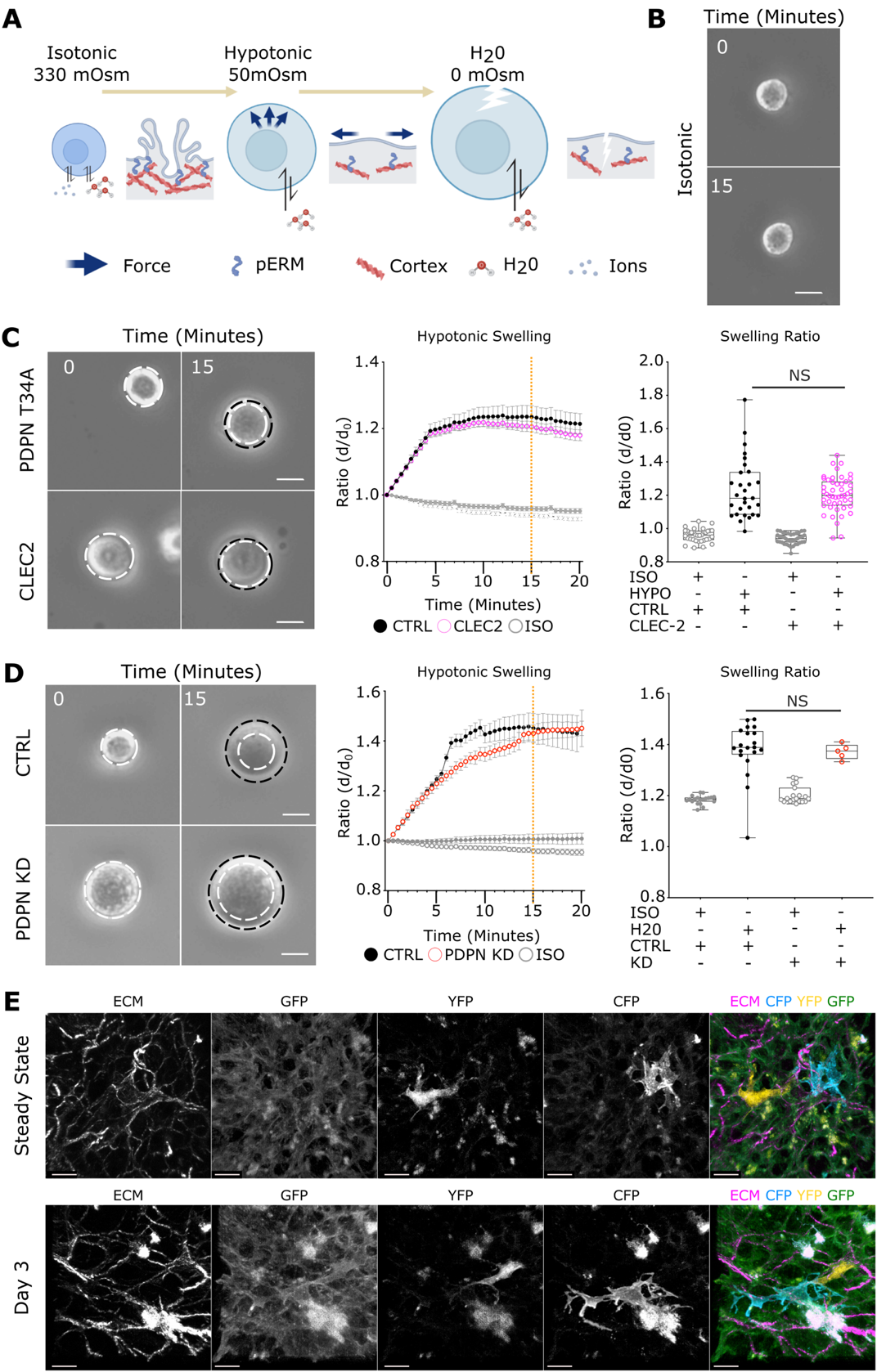
**Fig. S4 - Membrane access of fibroblastic reticular cells is increased through the CLEC2/PDPN signalling axis**

**A)** Schematic of osmotic shock experiment, inducing swelling by altering osmolarity. **B)** Representative stills of FRC in isotonic control media. Scale bar 25$\mu$m. **C)** Time course of PDPN T34A FRCs with or without CLEC2 treatment in hypotonic conditions. Stills (top) show swelling of cells in hypotonic media. White dotted circle marks initial size before swelling and is compared to swelling at t=20 (black dotted circle). Scale bar 25$\mu$m. Change in diameter ratio over time (bottom left, mean ±SEM). Swelling ratio comparisons between control and CLEC2 treatment at 15 minutes post swelling (orange dotted line, bottom right). Box plots indicates median and interquartile range. One-way ANOVA with Tukeys multiple comparisons. N>29. **D)** Time course of PDPN CTRL vs PDPN shRNA KD FRCs in extreme hypotonic conditions (H_2_0). Stills (top) show swelling of cells in hypotonic media. White dotted circle marks initial size before swelling and is compared to swelling at t=20 (black dotted circle). Scale bar 25$\mu$m. Change in diameter ratio over time (bottom left, mean ±SEM). Swelling ratio between control and PDPN shRNA KD FRCs at 15 minutes post swelling (orange dotted line, bottom right). Box plots indicates median and interquartile range. One-way ANOVA with Tukeys multiple comparisons. N>16. **E)** Single channel stills of PDGFR$\alpha$-mGFP confetti FRC network in the steady state and day 3 post immunisation. Scale bar 10$\mu$m.

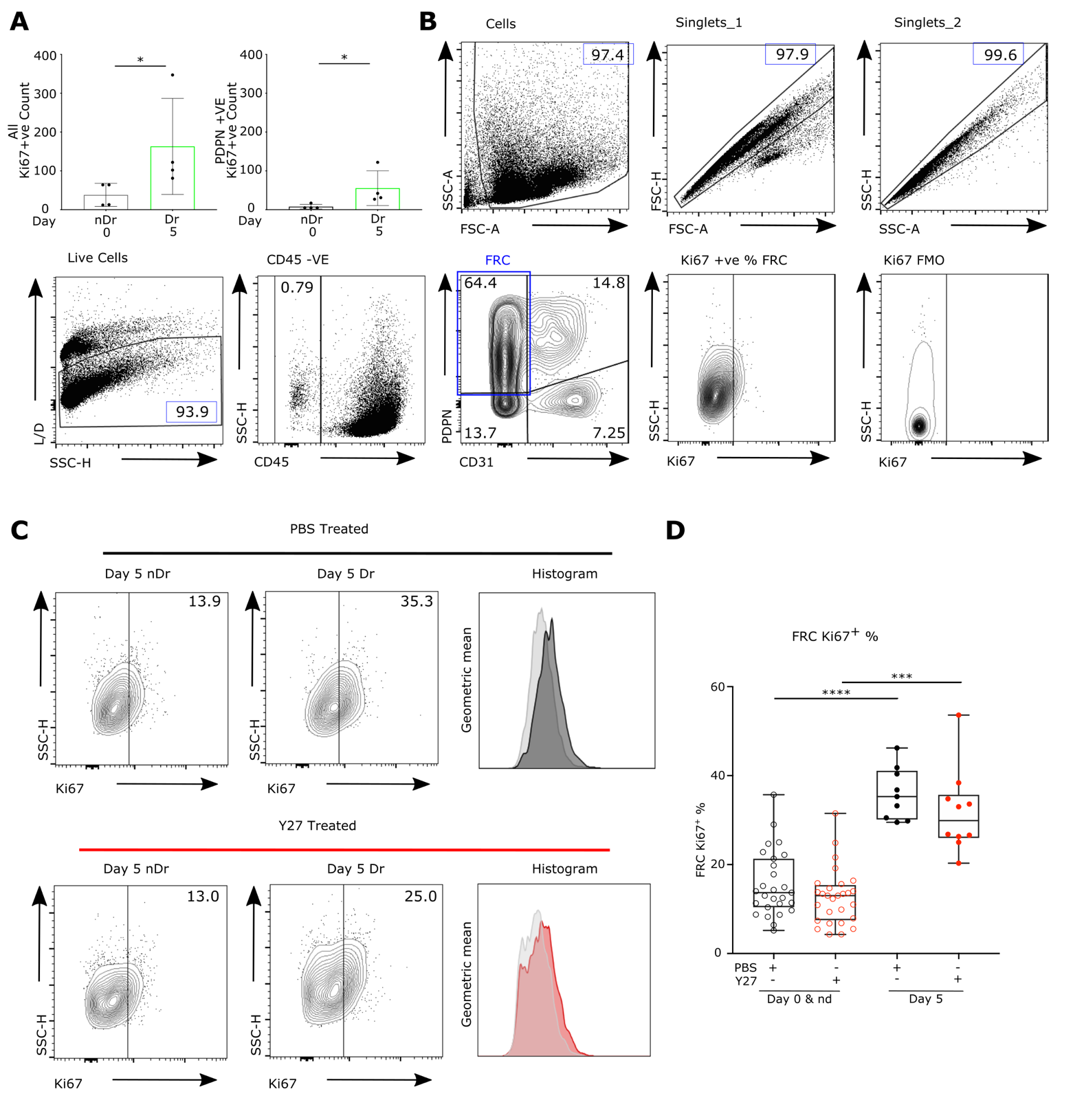

**Fig. S5** - **Flow cytometry gating strategy and Ki67 measurements**

**A)** Quantification of Ki67^+^ cells (top) and PDPN^+^ Ki67^+^ cells (bottom) per LN region. Box plots indicates median and interquartile range. Mann-Whitney test, p<0.05. N=2. **B)** Schema of flow cytometry gating strategy for measurement of stromal cell populations. **C)** Example of Ki67^+^ FRCs cell plots and histogram of Ki67 geometric mean, comparing immunisation and Y27 treatments. **D)** Quantification of Ki67^+^ FRCs. Box plots indicates median and interquartile range. Two-way ANOVA with Tukey’s multiple comparisons. ****p<0.0001, ***p<0.001. N>9 animals per condition

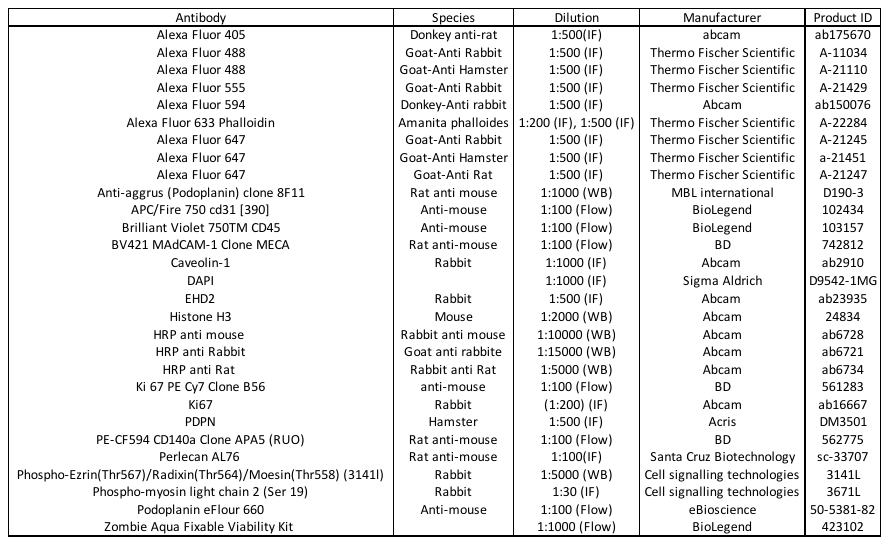
**Tables S1**

*Table S1* – **Antibody reagents and dilutions**

**Movies S1 to S4**

**S1A – The paracortical T-cell FRC network, showing 3D localisation of actomyosin structures in homeostasis**.

Staining of the conduit (perlecan, magenta), FRC network (PDPN, yellow), F-actin (phalloidin, cyan) and pMLC (red). PDPN and perlecan surface renders show the location of actomyosin in relation to these structures.
